# Samovar: Single-sample mosaic SNV calling with linked reads

**DOI:** 10.1101/560532

**Authors:** Charlotte A. Darby, James R. Fitch, Patrick J. Brennan, Benjamin J. Kelly, Natalie Bir, Vincent Magrini, Jeffrey Leonard, Catherine E. Cottrell, Julie M. Gastier-Foster, Richard K. Wilson, Elaine R. Mardis, Peter White, Ben Langmead, Michael C. Schatz

## Abstract

We present Samovar, a mosaic single-nucleotide variant (SNV) caller for linked-read whole-genome shotgun sequencing data. Samovar scores candidate sites using a random forest model trained using the input dataset that considers read quality, phasing, and linked-read characteristics. We show Samovar calls mosaic SNVs within a single sample with accuracy comparable to what previously required trios or matched tumor/normal pairs and outperform single-sample mosaic variant callers at MAF 5%-50% with at least 30x coverage. Furthermore, we use Samovar to find somatic variants in whole genome sequencing of both tumor and normal from 13 pediatric cancer cases that can be corroborated with high recall with whole exome sequencing. Samovar is available open-source at https://github.com/cdarby/samovar under the MIT license.

## 1 Background

Genomic mosaicism results from postzygotic *de novo* mutations, ranging from single-nucleotide changes to larger structural variants and whole chromosome aneuploidy. Mosaic mutations are present in some of the cells belonging to the offspring, but in none of either parents’ cells [1, 2]. The distribution and prevalence of cells with a mosaic mutation depend on a combination of the developmental cell lineage, stage at which the mutation occurred, selection for or against cells with the mutation [3], and cell migration [4]. Somatic mosaicism refers to genetic heterogeneity among non-germ cells, which accrue in normally dividing cells throughout the human lifetime [5, 6, 7] corroborated by monozygotic twin studies [8]. Mosaicism also plays an important role in many genetic diseases. Pathologically, cancer is characterized by an overall increased mutational load in tumor cells as well as a high level of intra-tumor genetic heterogeneity [9, 10]. Mosaicism has also been implicated in autism [11] and is being explored in connection to other neurological disease [12, 13, 14]. Causal mosiac mutations have also been found for Sturge-Weber syndrome [15], McCune-Albright syndrome [16] and Proteus syndrome [17] among others.

Mosaic variants can be detected by whole-genome or targeted sequencing of affected tissue. Samovar operates on linked reads, which are sets of sequencing reads deriving from a longer fragment such as those from the 10X Genomics Chromium instrument (Pleasanton, CA). While the individual (“constituent”) reads are typical short Illumina reads, the longer fragments can be tens or hundreds of kilobases long. The mapping from constituent reads to fragments of origin is established by molecular barcodes added in the Chromium library preparation step. The average sequencing coverage per long fragment is usually low: around 0.1-fold [18]. Since constituent reads can be paired-end, we use the term “long fragment” for the longer fragment from which a linked read is derived, and “short fragment” for fragments from which paired-end reads are derived.

The properties of linked reads enable many potential improvements in variant detection and related analyses [19]. For example, a constituent read that would align repetitively by itself might align uniquely when alignments of other reads from the same long fragment are accounted for [20, 21]. Linked read based algorithms have been developed for *de novo* assembly [22, 23, 24] *de novo* mutation calling [25], assembly error-correction [26] and structural variant calling [27, 28, 29, 30, 31]. Also, linked reads enable more accurate and contiguous assembly of haplotypes [18, 32] since constituent reads can be phased even when only some overlap heterozygous variants (Figure 1b).

**Figure 1:**
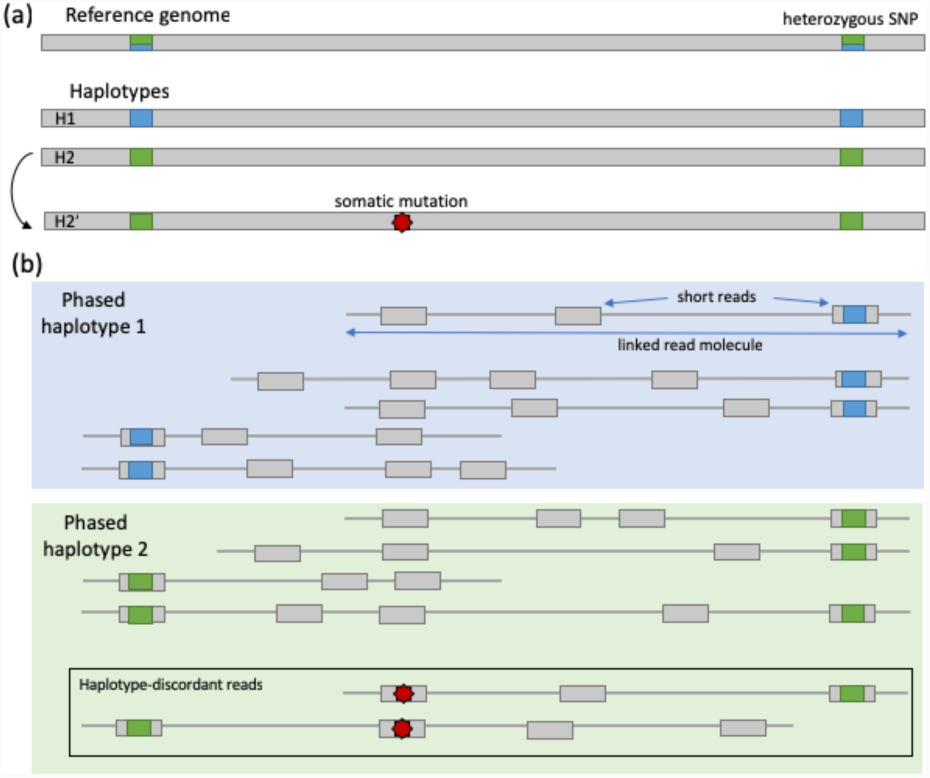
(a) A mosaic mutation occurs on haplotype H2. (b) Therefore, in linked read sequencing, where short reads can be phased when linked reads overlap phased heterozygous variants, mosaic mutations manifest on reads from only one haplotype, here H2. Adapted from Figure 3 of [33].

Though downstream tools benefit automatically from some linked-read properties — e.g. improved alignment accuracy — other benefits require specialized methods to exploit. In particular, the detection of a somatic mosaic SNV can benefit from haploytype assembly only if the model is informed by the mapping between constituent reads and linked reads. As an example, in a diploid sample with haplotypes H1 and H2, suppose a mosaic mutation occurs on haplotype H2 yielding a collection of reads (labeled H2’) that have the mosaic allele but otherwise match H2 (Figure 1a). The mosaic mutation will likely be tolerated by the haplotype assembler and the reads will still be assigned to H2 (Figure 1b). The fact that all the mosaic-carrying reads fall on the same haplotype is a hallmark of post-zygotic mosaicism [11] and contrasts with sequencing error, which would tend to distribute the “mosaic” alleles evenly across haplotypes [34]. Reads with the mosaic allele are called haplotype-discordant reads, and these are the most reliable kind of evidence we can gather in support of mosaic variants.

The mosaic variant caller’s task is to distinguish the signature of a mosaic variant from that of a germline variant after it has been affected by sequencing errors, alignment errors, copy-number changes and other confounders. Most methods employ statistical tests on the sequencing reads aligned to a particular site, comparing allele frequency between “tumor” and “normal” (or between the observed and expected value for a germline variant). See [33] for a review of methods to detect such mutations in scenarios other than cancer, and [35] for a comparison of several tools in the cancer context. Samovar is unique in that it is the first to evaluate haplotype-discordant reads identified through linked read sequencing, thus enabling phasing and moasic variant detection across essentially the entire genome. It also evaluates the statistical characteristics of the haplotypes, depth of coverage, and potential confounders such as alignment errors, to robustly identify mosaic variants from a single sample.

## 2 Results

### 2.1 Samovar pipeline

We present Samovar, a single sample mosaic SNV caller designed for 10x Genomics linked-read whole-genome sequencing (WGS) data. Samovar takes as input phased variants in VCF format and linked-read alignments in BAM format. These are both output by 10x Genomics’ Long Ranger pipeline, which preprocesses reads, aligns linked reads, calls variants and assembles haplotypes.

The Samovar workflow is shown in Figure 2, and proceeds in six major steps. In step 1, Samovar identifies all genomic sites where there is sufficient data to apply our model. This is done by filtering based on features such as depth of coverage, fraction of reads that are phased, frequency of the candidate mosaic allele, and related data characteristics. In step 2, Samovar modifies the input BAM file to introduce synthetic mosaic variants to be used as sample-specific training data. Specifically, these variants are used as positive examples for training our model, whereas real homozygous/heterozygous variants, as called by Long Ranger, are used as negative examples. In step 3, Samovar trains a random forest model containing an ensemble of 100 individual decision trees that scores sites according to their resemblance to the synthetic-mosaic sites. In step 4, Samovar scores all sites that passed the initial filter using this model. In step 5, complex repeat regions and non-diploid copy-number regions are optionally filtered out. In step 6, a final filter removes false positives resulting from alignment errors to produce scored mosaic variant calls.

**Figure 2:**
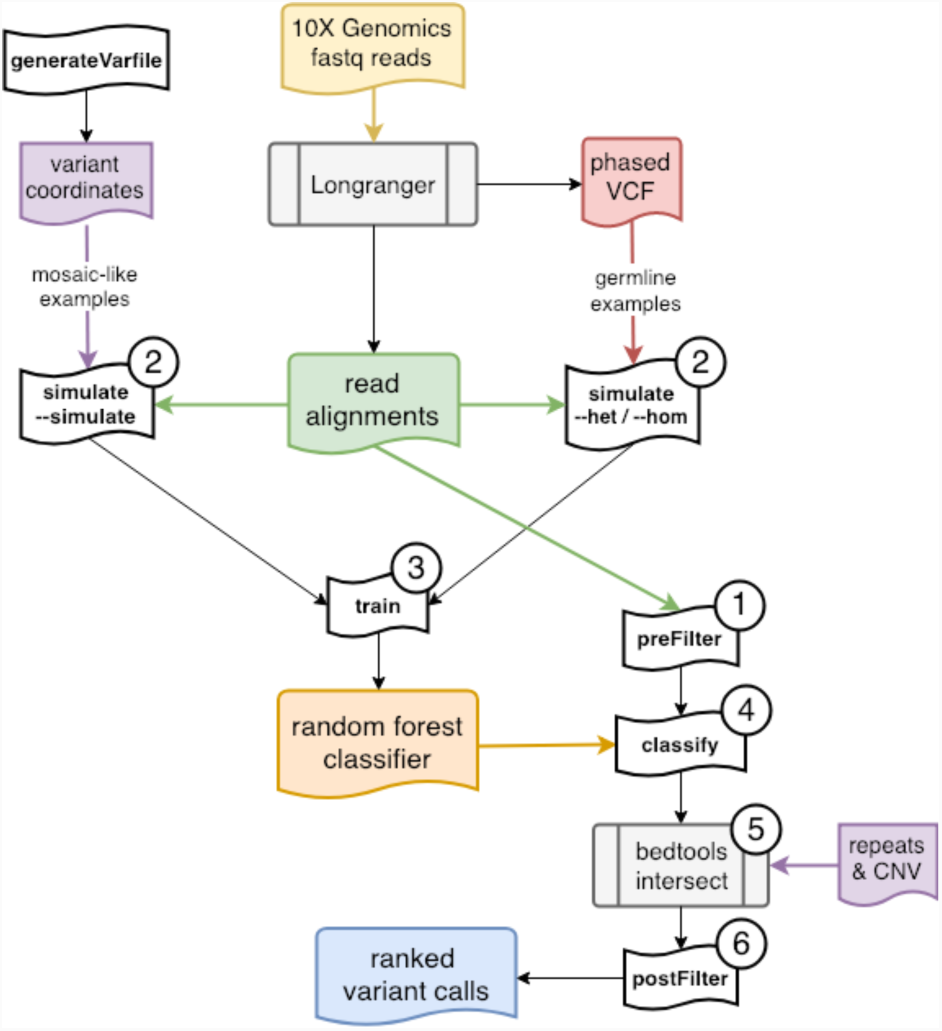
Samovar workflow.

### 2.2 Simulation experiment

To benchmark Samovar, we used bamsurgeon [36] to insert synthetic mosaic variants into the NA24385 10x Genomics Chromium BAM file from the Genome in a Bottle project [37]. Training and testing occurred using sites on the autosomal chromosomes only since NA24385 is male. The mean inferred linked read length is 16,176 bp with standard deviation 54,387 bp. To evaluate performance at lower coverage and in other tools’ paired mode, the original BAM file (mean coverage 61.8; median 60 at bamsurgeon-modified sites, excluding reads marked duplicate) was split in half based on read group tag and we subsequently modified only one half with bamsurgeon (mean coverage 30.6, median 29 at bamsurgeon-modified sites). Splitting by read group tag ensures that an entire linked read will be placed into the derivative BAM file. Experiments with the original BAM file are referred to as “60X coverage” and those with the subsample as “30X coverage.”

#### Samovar model comparison

To measure the specific advantage conferred by linked reads, we also implemented two reduced Samovar models that incorporate less of the variant phasing information. The “short-only” model redefines the fragment-level model features so that they use information summarized over the shorter, paired-end-level fragments rather than the longer linked-read-level fragments. In this model, a paired-end read is assigned to a haplotype only if one of the ends overlaps a heterozygous variant phased by Long Ranger. Past work showed that even the phasing information from short fragments can improve mosaic variant calling accuracy [34]. We find while the precision is comparable to the Samovar full model, the number of variant calls is much lower, resulting in a genome-wide recall of 2.0% at 30X and 60X, because there are few sites for which adequate phasing information can be compiled from short reads alone (Figure S4, Table S4).

We also created a “no-phasing” Samovar model that used no fragment phasing information at all. This was accomplished simply by omitting the fragment-level features from the model. Precision in every MAF bin is near zero, although genome-wide recall is 68.3%, underscoring the importance of phasing features to our approach (Figure S4, Table S4).

#### MosaicHunter and MuTect2 comparison

We compared Samovar to MosaicHunter v. 1.1. [38]. We ran MosaicHunter in “tumor-only mode” analyzing only the bamsurgeon-mutated BAM file from NA24385, as well as in “trio mode” where the unaltered GIAB 10x Genomics Chromium BAM files from the mother (NA24143) and father (NA24149) were also provided. The parental BAM files were similarly produced by Long Ranger but not modified by bamsurgeon. While Samovar does not use trio information, we hypothesized that its modeling of linked-reads would allow it to have competitive accuracy. The modified and unmodified halves of the BAM file split by read group were provided when MosaicHunter was run in “paired-mode” as tumor and normal, respectively.

We also compared Samovar to MuTect2 from GATK v. 4.0.12.0. [39]. We ran MuTect2 in “tumor-only mode” and “paired-mode” on the same data described above. Tumor-only mode calls mosaic and germline mutations simultaneously but does not differentiate between the categories; hence the number of calls is much higher.

Figure 3 shows each tool’s precision, stratified by MAF in the tumor WGS. Precision is calculated as the fraction of variant calls made that were bamsurgeon synthetic mutations. Samovar achieves consistently higher precision than the tumor-only modes of MuTect2 and MosaicHunter. Importantly, Samovar’s precision is also comparable to those tools in their trio and paired modes, with MosaicHunter’s paired and trio modes achieving slightly higher precision at MAFs ≥ 0.2 and MuTect2’s paired mode achieving higher precision at MAFs ≥ 0.3.

**Figure 3:**
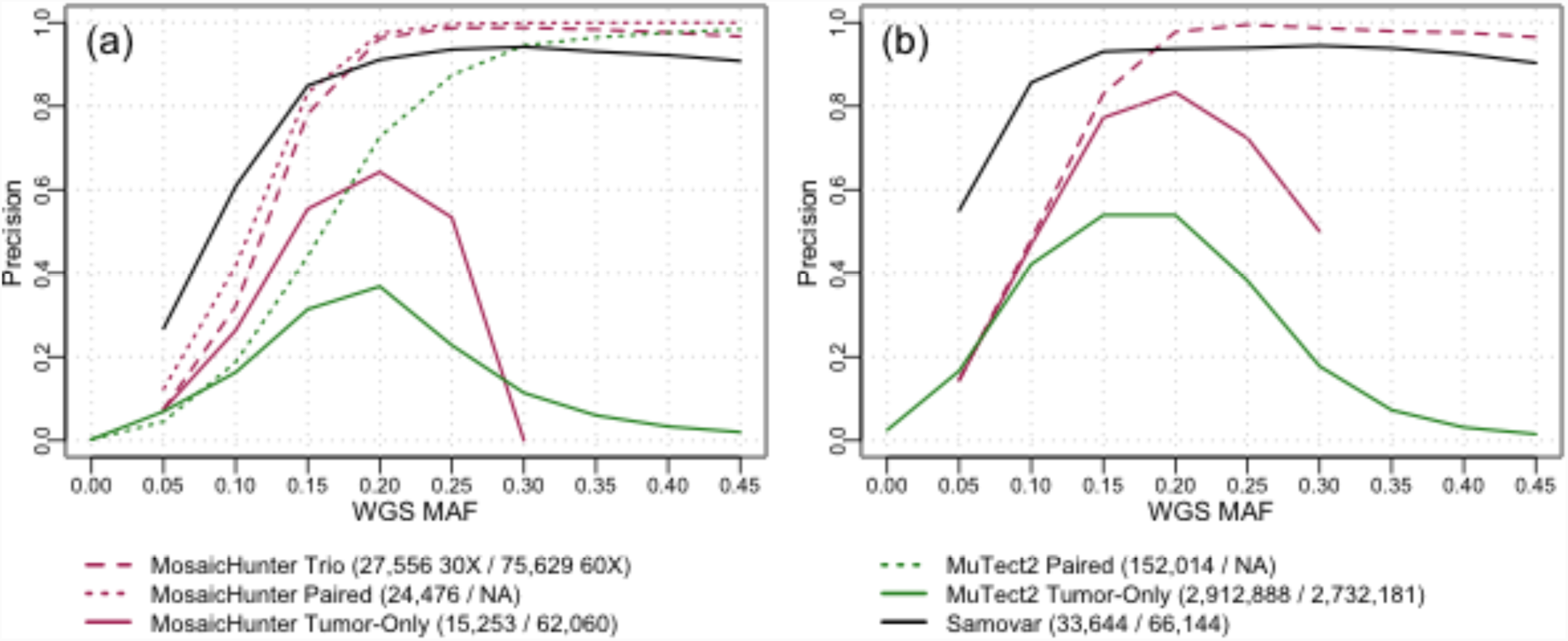
Precision calculated for Samovar, MuTect2, and MosaicHunter variant calls stratified by mosaic allele fraction (MAF) in the whole genome sequencing data (WGS). (a) 30X coverage (b) 60X coverage.

Note that in all cases, the original 10x Genomics BAM file was used. This means that all three Samovar models (as well as MuTect2 and MosaicHunter) benefited from the improved alignment accuracy of the linked-read-aware Lariat aligner, giving the short-only and no-phasing models and the other two methods a somewhat artificial advantage.

In addition to performance genome-wide we evaluated precision and recall (i.e. TPR) across different annotated genomic regions: genes, exons, all repeats, Alu repeats, segmental duplications, enhancers and promoters listed in the UCSC Genome Browser and Ensembl, shown in Table 1. Recall is calculated as the fraction of bamsurgeon synthetic mutations with at least four mosaic allele reads that were in the variant call set since both Samovar and MosaicHunter require at least four reads to support a variant call. In practice, many tools including Samovar and MosaicHunter apply filters that exclude portions of the genome that lack sufficient evidence or that are inherently difficult to analyze, such as highly repetitive portions, which particularly contributes to MosaicHunter’s poor performance in these genomic regions. (See Note S4.) Furthermore, 66% of the Samovar false negative sites over which recall was evaluated in the 30X coverage experiment and 38% of false negatives in the 60X experiment had fewer than four haplotype-discordant reads, which is the default requirement for Samovar. Relaxing this parameter can boost recall, although may also impact precision.

**Table 1:** Precision (Prec), recall (Rec), and F_0.5_ score (F measure with β = 0.5) of each tool for the synthetic mosaic variants inserted by bamsurgeon.

| 30X Coverage | Samovar |  |  | MuTect2 |  |  |  |  |  | MosaicHunter |  |  |  |  |  |  |  |  |
| --- | --- | --- | --- | --- | --- | --- | --- | --- | --- | --- | --- | --- | --- | --- | --- | --- | --- | --- |
|  |  |  |  | Tumor-Only |  |  | Paired |  |  | Tumor-Only |  |  | Paired |  |  | Trio |  |  |
| | Prec | Rec | $F_{0.5}$ | Prec | Rec | $F_{0.5}$ | Prec | Rec | $F_{0.5}$ | Prec | Rec | $F_{0.5}$ | Prec | Rec | $F_{0.5}$ | Prec | Rec | $F_{0.5}$ |
| Autosomes | 84.0 | 30.1 | 61.9 | 3.0 | 82.3 | 3.7 | 60.8 | 91.4 | 65.1 | 31.5 | 5.1 | 15.5 | 79.2 | 20.7 | 50.5 | 70.4 | 20.7 | 47.5 |
| Exons | 84.0 | 28.3 | 60.3 | 3.6 | 85.5 | 4.5 | 60.1 | 92.0 | 64.6 | 35.0 | 7.1 | 19.6 | 82.1 | 30.8 | 61.6 | 73.7 | 30.8 | 57.6 |
| Genes | 84.9 | 30.1 | 62.2 | 3.2 | 83.5 | 4.0 | 63.0 | 92.0 | 67.3 | 32.6 | 5.7 | 16.7 | 79.9 | 22.7 | 53.2 | 71.2 | 22.7 | 49.9 |
| Enhancer | 88.5 | 31.0 | 64.6 | 3.9 | 84.6 | 4.9 | 72.9 | 92.3 | 76.1 | 37.8 | 5.9 | 18.1 | 85.5 | 29.5 | 62.0 | 80.2 | 29.5 | 59.7 |
| Promoter | 83.3 | 26.1 | 57.9 | 3.0 | 81.5 | 3.8 | 59.4 | 90.9 | 63.9 | 35.3 | 6.1 | 18.0 | 80.5 | 25.1 | 55.9 | 73.7 | 25.1 | 53.1 |
| Alu | 82.0 | 28.6 | 59.7 | 2.3 | 77.0 | 2.9 | 54.5 | 88.4 | 59.0 | 8.6 | 0.0 | 0.2 | 56.5 | 0.3 | 1.4 | 53.1 | 0.3 | 1.4 |
| RepeatMasker | 84.2 | 29.6 | 61.6 | 2.8 | 80.6 | 3.4 | 58.9 | 90.1 | 63.2 | 20.2 | 0.3 | 1.4 | 72.3 | 1.4 | 6.4 | 61.3 | 1.4 | 6.4 |
| Seg. Dup. | 25.6 | 10.4 | 19.8 | 1.3 | 55.7 | 1.6 | 18.4 | 62.8 | 21.4 | 6.6 | 0.5 | 1.8 | 39.3 | 1.7 | 7.1 | 29.1 | 1.7 | 6.8 |
| 60X Coverage | Prec | Rec | $F_{0.5}$ | Prec | Rec | $F_{0.5}$ | | | | Prec | Rec | $F_{0.5}$ | | | | Prec | Rec | $F_{0.5}$ |
| Autosomes | 84.6 | 43.0 | 70.9 | 3.6 | 76.0 | 4.5 |  |  |  | 32.4 | 15.5 | 26.6 |  |  |  | 46.8 | 27.2 | 40.9 |
| Exons | 84.3 | 41.8 | 70.1 | 4.7 | 79.6 | 5.8 |  |  |  | 38.5 | 25.3 | 34.9 |  |  |  | 54.0 | 45.5 | 52.1 |
| Genes | 85.6 | 43.4 | 71.7 | 3.9 | 77.2 | 4.9 |  |  |  | 33.1 | 17.0 | 27.8 |  |  |  | 47.7 | 30.0 | 42.6 |
| Enhancer | 90.8 | 47.8 | 77.0 | 4.8 | 77.9 | 5.9 |  |  |  | 36.9 | 22.7 | 32.8 |  |  |  | 51.6 | 40.0 | 48.8 |
| Promoter | 85.4 | 40.7 | 70.0 | 4.0 | 76.8 | 4.9 |  |  |  | 38.5 | 21.1 | 33.0 |  |  |  | 56.4 | 40.5 | 52.3 |
| Alu | 81.1 | 42.9 | 68.8 | 3.0 | 68.0 | 3.6 |  |  |  | 16.5 | 0.2 | 1.1 |  |  |  | 31.7 | 0.5 | 2.3 |
| RepeatMasker | 84.2 | 42.2 | 70.2 | 3.4 | 74.1 | 4.1 |  |  |  | 24.7 | 1.0 | 4.3 |  |  |  | 38.3 | 1.8 | 7.4 |
| Seg. Dup. | 28.0 | 13.1 | 22.8 | 1.6 | 48.5 | 2.0 |  |  |  | 9.8 | 1.5 | 4.7 |  |  |  | 18.5 | 2.7 | 8.6 |

### 2.3 Pediatric cancer

We next studied a collection of 13 pediatric cancer cases that we sequenced — both tumor and normal – using 10x Genomics Chromium Whole-Genome Sequencing (WGS) and Whole-Exome Sequencing (WES). One of these cases was studied previously [40], and the other twelve are novel to this work. We ran Samovar, MosaicHunter (in both paired and tumor-only modes), and MuTect2 (in both paired and tumor-only modes) on each of the 13 tumor WGS datasets. When running MosaicHunter or MuTect2 in paired mode, we also provided the paired normal WGS.

To estimate accuracy of the different approaches, we used the WES sequencing as a validation dataset as it provides independent and deeper coverage over candidate variants within the exome. We first identified the calls from each tool within the exome capture region. The number and precision of the exome-coincident calls made by each tool are shown in Table 2.

**Table 2:**
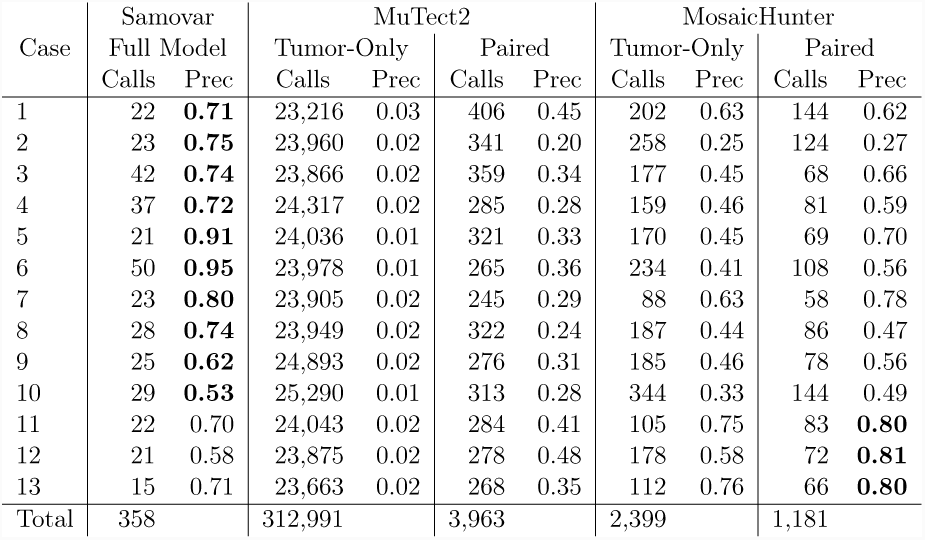
Number of variant calls in the exome capture regions and precision (Prec) based on supporting reads found in WES. Samovar has the highest validation rate in 10 out of the 13 cases.

We then examined the corresponding WES tumor data for evidence of the mosaic call made in the WGS data. We considered a mosaic variant call to be “validated” if (a) the corresponding WES tumor sample had at least 50 aligned reads at the locus with at least 4 reads supporting the mosaic allele, and (b) the mosaic variant was not found to be germline by Long Ranger in both the tumor and normal WGS data from that patient. Figure 4 stratifies the validation rate by MAF in the WGS data and Table 2 shows each tool’s overall precision for the calls in the exome capture region. The bar graph shows the number of variants in each MAF bin. MosaicHunter paired called 3 times as many variants as Samovar, and MuTect2 paired called 11 times as many variants. This is because Samovar requires phasing-based evidence to make a call. However, Samovar’s validation rate is comparable to the paired callers across a range of MAF, indicated by the comparable precision of Samovar in Figure 4e compared to other tools’ paired modes in a and c. Against tumor-only modes of other tools, Samovar has superior precision especially at MAF ≥ 0.15: MuTect2 tumor-only mode is not designed to differentiate heterozygous from high-MAF mosaic variants, and MosaicHunter makes few calls with a low validation rate.

**Figure 4:**
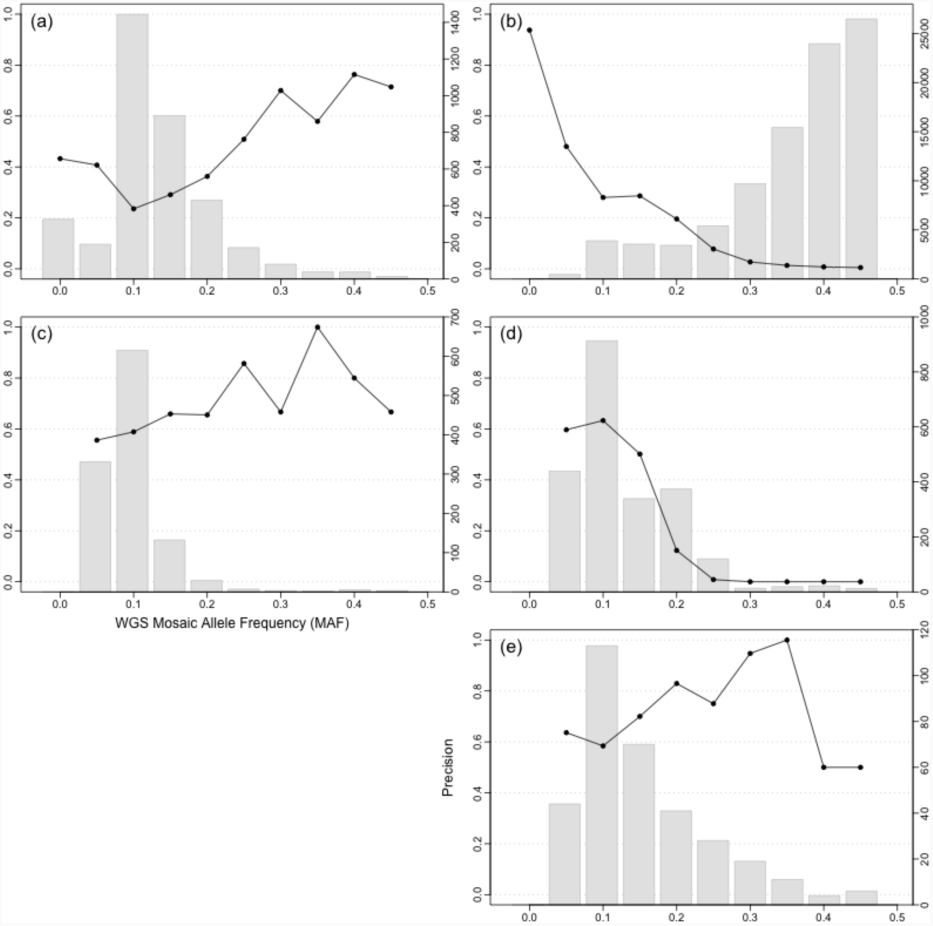
Fraction of variant calls in exome capture region supported by WES data (black line, left axis ticks) and number of variant calls (gray bars, right axis ticks) stratified by mosaic allele fraction (MAF), combined for the 13 pediatric cancer cases studied. (a) MuTect2 paired (b) MuTect2 tumoronly (c) MosaicHunter paired (d) MosaicHunter tumor-only (e) Samovar.

As Samovar demonstrated high single sample precision in simulation, comparable to the other tools’ paired analysis, we are also able to run it on the normal control available for each of these cases. Sensitivity was measured in the same fashion using WES of the normal sample; across all 13 samples, 732 variants were in the exome capture region and the validation rate was 65% (see Table S9 for per-sample statistics). Interestingly, using ANNOVAR [41], we determined 11 of these mosaic mutations across 7 cases were nonsynonymous (amino-acid-changing) in one of the 299 cancer driver genes listed in [42]. The extent of mosaicism in normal tissue and how this may relate to pediatric cancer are interesting avenues of future study now possible with Samovar.

## 3 Methods

### 3.1 Samovar pipeline

Samovar is implemented in Python 3 and operates on the alignment (BAM) and variant (VCF) files produced by 10x Genomics’ Long Ranger pipeline. See Note S1 for software dependency and input file requirements.

#### (1) preFilter

Samovar first scans the genome calculating the features listed in Figure S8 at each site. Each feature has a numerical threshold, and if all filters are passed the site is considered in step 4 (classify) as a candidate variant site. These filters examine measurements such as depth, number of haplotype-discordant reads, quality of the alignments and credibility of the read phasing.

#### (2) simulate

Simulated mosaic training examples are generated at regular intervals across the genome at a range of mosaic allele frequency (MAF) from 0.025 to 0.475 at increments of 0.025. Such sites are called “simulation sites.” Sites harboring germline variant calls can be excluded by specifying them in a VCF. For each phased alignment having the reference allele at the simulation site, the reference allele is randomly changed to the mosaic base with probability equal to the target MAF. For an unphased alignment having the reference allele, the reference allele is randomly changed to the mosaic base with probability 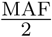, on the principle that unphased reads are equally likely to originate from either haplotype. The features listed in Figure S5 are computed for the simulation sites to obtain true-mosaic training examples. The same features are computed for FILTER=PASS phased heterozygous (GT=0|1 or GT=1|0) and homozygous (GT=1|1 or GT= 0|0) variant sites from the VCF to get true-non-mosaic examples.

#### (3) train

A random forest model is trained with an equal number of simulation sites and non-mosaic sites. Non-mosaic sites are selected to have equal amounts of heterozygous and homozygous calls in the VCF. We use the RandomForestClassifier module from the scikitlearn library [43] with `max_leaf_nodes 50` and `n_estimators 100`, though Samovar allows the user to customize these hyperparameters. The random forest features described in Table S5 take into account the abundance and consistency of evidence for a mosaic variant, including the number of haplotype discordant reads, mosaic allele fraction, base quality, alignment score, amount of soft clipping, presence of indels, etc.

After cross-validation at a variety of sequencing depths (Table S1), we found that using 20,000 mosaic, 10,000 heterozygous and 10,000 homozygous training examples achieved a balance of computational efficiency and accuracy. We subsampled the NA24385 BAM file used for the simulation experiment and ran the Samovar simulate and train steps. For each number of training examples, average performance statistics are reported for ten independent train/validation splits; 0.5 and 0.9 refer to the random forest probability that the example is in the mosaic class.

#### (4) classify

Genomic sites passing the preFilter are classified by the trained random forest model, yielding the predicted probability that the site is mosaic. Sites with probability above a cutoff are reported in BED format. Based on cross-validation at a variety of sequencing depths, we found that a probability cutoff of 0.5 balances false positive rate and true positive rate (Table S1), though this can be adjusted to trade between sensitivity and precision.

#### (5) region-based filter

As Illumina sequencing is known to have high error rates within microsatellites and simple repeat sequences [44], we exclude candidate mosaic variants idenfied in these regions. Specifically, we exclude variants within +/-2bp from 1,2,3,4-bp repeats at least 4bp long with at least 3 copies of the unit. Within hg19, 72.0% of autosomes and 71.4% of autosomes+X+Y will remain after this region filter, and within GRCh38 73.8% of autosomes and 73.1% of autosomes+X+Y remain. We also exclude any CNV regions +/-5bp identified by CNVNATOR [45] because polymorphism among the copies of a repeated region would be misconstrued as mosaicism.

#### (6) postFilter

Our expectation is that mosaic variants are isolated events. Samovar applies a final test to distinguish an isolated, likely mosaic variant from the situation where there are many nearby variants co-occuring on the same reads. The latter pattern is usually caused by alignment errors in the presence of repetitive DNA and copy number variation. Specifically, we examine each base within a fixed distance of the mutative mosaic locus. At each base we conduct a Fisher’s exact test, testing if the alleles observed at the query base associate with the haplotype-discordant reads. If the most significant p-value among all the statistical tests is less than the threshold, the site is filtered out. Based on simulations, we find that the p-value threshold can be set to 0.005 (default) or lower based on the desired balance between precision and recall. There is an option to avoid particular sites when calculating the minimum p-value among all nearby sites and it is recommended to use the germline VCF of variant calls here.

The final mosaic variant calls are reported in VCF format. VCF INFO tags are used to convey information such as the number of haplotype-discordant reads, the model-predicted probability, and the minimal p-value obtained by the postFilter.

### 3.2 Simulation experiment

#### Input data

We downloaded the 10x Genomics Chromium datasets for the A/J trio processed with Long Ranger version 2.1 and GRCh38 from the GIAB project: FTP Link We use the BAM file from sequencing the son’s genome (NA24385) as the basis for this simulation experiment, but MosaicHunter uses the BAM files for the mother (NA24143) and father (NA24149) in trio mode.

We use a custom fork of bamsurgeon [36] to edit the reads in the BAM file. Given a target MAF, a 2 × MAF fraction of reads with tag HP=1, and a MAF fraction of reads with no HP tag are selected to mutate. The alternate allele is chosen randomly among the three non-reference bases.

Simulated mosaic mutations were introduced at evenly spaced intervals every 20,000 bp on the autosomes with target MAF between 0.025 and 0.475 in increments of 0.025. Reads were realigned with BWA-MEM after mutations were introduced. To compute precision, the denominator is sites with at least 4 alt-allele reads and 16 total reads (not marked duplicate or QC fail). This is because the parameters we chose for Samovar and MosaicHunter require at least 4 reads to call a mosaic variant, and Samovar’s depth filter threshold is 16 (MosaicHunter’s minimum depth is 25, which we keep, so technically fewer sites are visible to MosaicHunter).

#### Samovar

We use 20,000 simulated mosaic, 10,000 heterozygous and 10,000 homozygous training examples to train each random forest model described. Table S3 has the feature importances of the Samovar model, with abbreviation and number as in Figure S5.

#### Samovar Short-read phasing model

Samovar is designed to take advantage of the long-range phasing information given by linked reads. Previous methods similarly took advantage of the shorter-range phasing information given by paired-end sequencing. We can simulate the paired-end strategy in Samovar, allowing us to compare to the linked-read strategy while holding the rest of the pipeline constant. We begin by creating a “short-read phasing” Samovar model that breaks down the linked reads into their constituent paired-end reads and considers only these shorter fragments when compiling linked-read-related features such as haplotype-discordant reads.

Supposing that we have the complete haplotype phasing from Long Ranger, we assign a haplotype to a pair of reads if either mate overlaps at least SNP with a phased genotype in the VCF. Out of 1.91 billion reads, 9.76% of reads could be phased. Only 0.006% of reads overlapped variants but had alleles for conflicting haplotypes – these were not phased. Table S6 has the feature importances of this limited model, with abbreviation and number as in Figure S6.

#### Samovar No-phasing model

While we do not advocate this approach, for the purposes of comparison, we remove all phasing-related features from Samovar to create a “no-phasing” model. Table S7 has the feature importances of this limited no-phasing model, with abbreviation and number as in Figure S7. Filters use the default parameters described in the preFilter feature list (Figure S8).

#### MosaicHunter

Version 1.1. We used the default recommended parameters when possible, except we did not use the misaligned_reads_filter because it was extremely slow. In addition, because we have simulated far more mosaic sites than would be expected in a normal genome, we do not want to penalize MosaicHunter because it deliberately filters mosaic sites that are close to each other so we changed the following parameters:

- `clustered_filter.inner_distance=2000` [default 20000]
- `clustered_filter.outer_distance=2000` [default 20000]

We also adjusted MosaicHunter’s supporting read threshold since Samovar requires at least 4 minor (mosaic) allele reads using: `base_number_filter.min_minor_allele_number=4` [default 3]

We used liftOver to transfer the provided `WGS.error_prone.b37.bed` and `all_repeats.b37.bed` to GRCh38 coordinates, and downloaded `dbsnp_human_9606_b150_GRCh38p7` bed files for the `common_site_filter`, `repetitive_region_filter`, `mosaic_filter.dbsnp_file` respectively.

Note that the `homopolymers_filter`, `common_site` and `repetitive_region` BED files leave visible only 32.2% of bases in the GRCh38 autosomes (34.4% including X and Y) to call mosaic variants. For comparison, Samovar considers about 73% of GRCh38 visible.

#### MuTect2

Version 4.0.12.0. We executed the standard GATK workflow of the Mutect2 program followed by FilterMutectCalls.

#### Genomic feature analysis

knownGene, knownGene exons, RepeatMasker, RepeatMasker Alu, Segmental duplications are from UCSC Table Browser (GRCh38, accessed 10/02/18). Ensembl Enhancer, Ensembl Promoter + flanking are from Release 94. Ensembl FTP Site

### 3.3 Pediatric cancer

#### Genomic DNA samples

Peripheral blood and paired tumor samples were obtained from patients enrolled onto the “Nationwide Children’s Neuro-Oncology Tumor and Epilepsy Tissue Bank” protocol (IRB16-00777) at Nationwide Children’s Hospital. 13 cases with paired blood and tumor derived DNA were extracted following the manufacturers recommendation using the AllPrep Kit for tumors (Qiagen) and Gentra Purgene or QIAamp Kit (Qiagen) for blood samples. Genomic DNA was quantified with the Qubit dsDNA HS Assay Kit (Life Technologies) and diluted to approximately 1 ng/*μ*L final concentration. DNA source and input mass into sample preparation is described in Supplementary File 1.

#### Sample preparation and sequencing

Linked-read whole genome sequencing (WGS) and whole exome sequencing (WES) libraries were generated [23]. Partitioning and barcoding high molecular weight (HMW) DNA was performed using a Chromium Controller Instrument (10x Genomics, CA), and Illumina sequencing libraries were prepared following protocols described in the manufacturer’s user guide (Chromium Genome Reagent Kits v2 -Rev A). For WES, 250 ng of each 10x linked-reads library was hybridized in pools (see Supplementary File 1) with 3 pmol of the xGEN Exome Research Panel v1.0 (Integrated DNA Technologies, Coralville, IA) per the manufacturers protocol. Post WES enrichment used standard Illumina P5 and P7 primers [46], and PCR cycling is highlighted in Supplementary File 1. Final libraries were quantified by qPCR (KAPA Biosystems Library Quantification Kit for Illumina platforms), diluted to 3 nM and sequenced using a paired-end recipe on the Illumina HiSeq 4000 next-generation sequencing instrument.

#### Bioinformatic Analysis

Cases using reference genome GRCh38 2.1.0 (1, 2, 7, 10, 11) were processed with Long Ranger 2.1.6 and GATK HaplotypeCaller 3.8-0. Samples using reference genome b37 2.1.0 (3, 4, 5, 6, 8, 9, 10, 12) were processed with Long Ranger 2.1.3 and GATK HaplotypeCaller 3.5-0. The sequencing coverage and fraction of the genome identified by the CNVNATOR [45] calls is recorded in Table S2, and the oncology diagnosis of each case in Table S8.

### 3.4 Computational efficiency

We report timing results for the 30X GIAB sample. Samovar completed in 7 hours with 48 parallel threads for the “filter” step and up to 4 parallel threads for other steps. MuTect2 paired mode completed in 136 hours with 48 parallel threads. MosaicHunter tumor-only and trio modes completed in 29 hours each and paired mode completed in 7 hours. Note MosaicHunter does not offer parallelism options. (See Note S3 for details.)

## 4 Discussion

Genomic mosaicism is an important characteristic of many human diseases and conditions. Accurately identifying mosaic variants has previously relied on paired samples or trio analysis, which increase study costs and complexity of studies and may not be possible in some situations. By taking advantage of linked-read properties — particularly the ability to accurately assemble haplotypes — Samovar is able to call mosaic SNVs for a single sample at a level of precision that is comparable to paired and trio-based methods. Samovar also achieves substantially higher precision at low MAFs (*<* 15%) and higher recall in more difficult-to-analyze portions of the genome such as segmental duplications and repetitive elements. This opens the door to a wider range of discoveries than are possible with current methods.

Though Samovar already compares favorably to tools that use matched-normal and trio data, in the future it will be important to investigate whether Samovar’s recall and precision can be further improved by incorporating trio and matched-normal data directly into it’s model. Based on the results collected here, we expect that a key benefit of this would be to improve recall at all MAFs and to extend the high precision achieved by the existing paired-and trio-based methods into the low end of the MAF spectrum.

## 5 Funding & Acknowledgements

This work was supported by the Nationwide Children’s Hospital Foundation. It was also supported by NIH grants U01MH106884 to BL, R01GM118568 to BL, R01-HG006677 to MCS, and R21-CA220411 to MCS; and NSF grant DBI-1350041 to MCS. Part of this research project was conducted using computational resources at the Maryland Advanced Research Computing Center (MARCC).

## Data Availability

The 10X Genomics linked read whole genome sequencing (WGS) and whole exome sequencing (WES) data described for the thirteen pediatric cancer cases is in the process of being deposited in dbGaP. The GIAB BAM files with simulated mutations are available at http://share.schatz-lab.org/samovar/simulation.

