## Supplemental Materials for "Samovar: Single-sample mosaic SNV calling with linked reads"

### Supplemental Materials for "Samovar: Single-sample mosaic SNV calling with linked reads" by Darby et al.

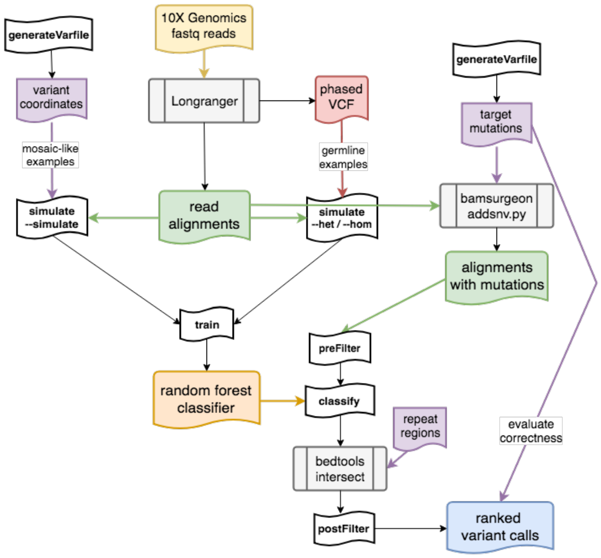

*Figure S1: Simulation experiment workflow (left) additionally evaluates correctness of the calls based on mutations generated with bamsurgeon.*

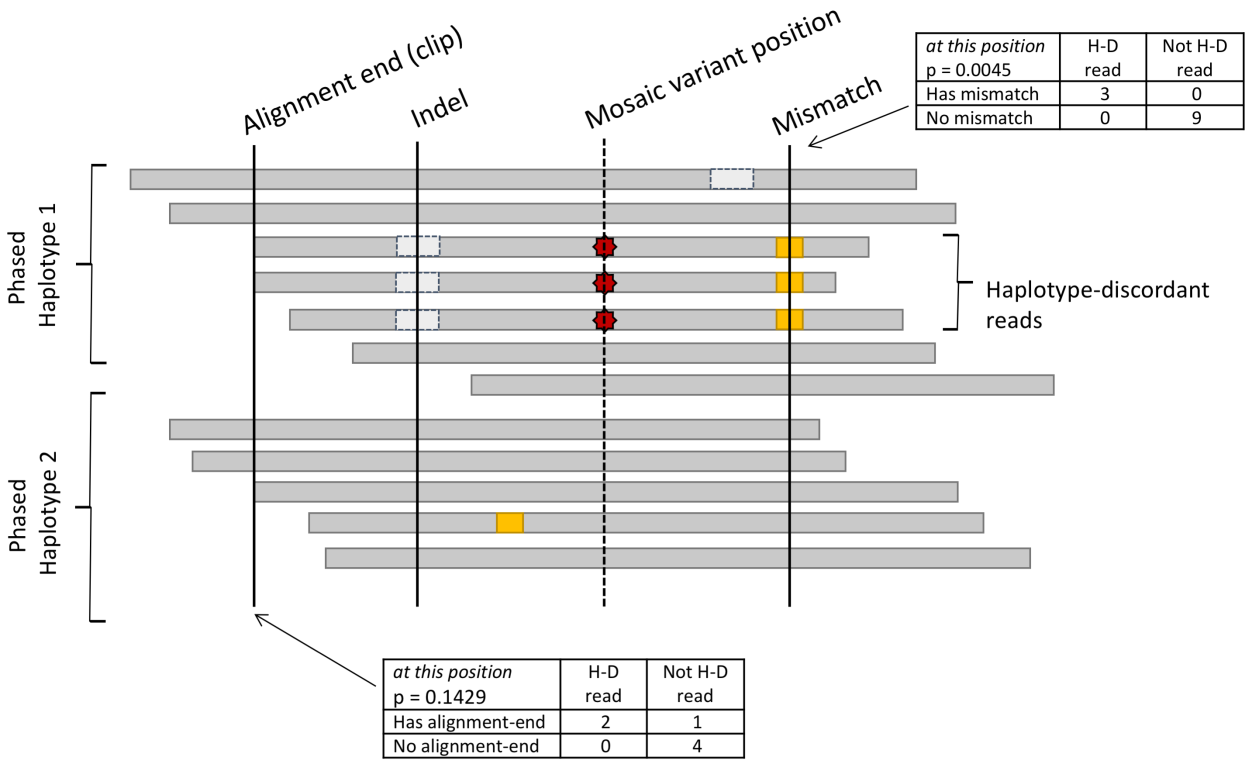

*Figure S2: The postFilter step calculates statistical association between haplotype-discordant reads and alignment features such as start/end position, indel or mismatch.*

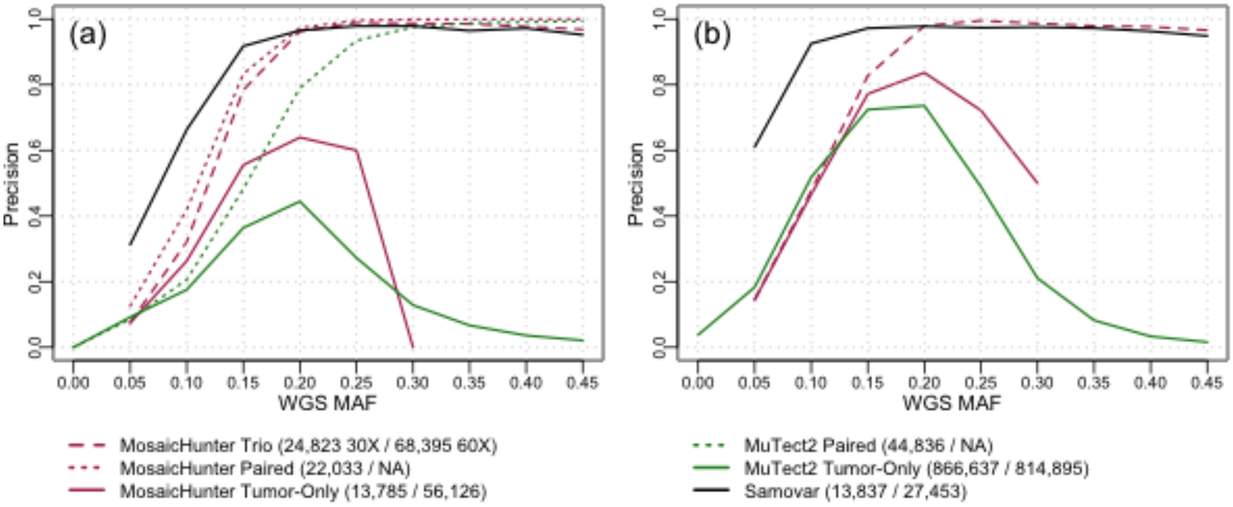

*Figure S3: Precision calculated in the genomic region not filtered by MosaicHunter or Samovar's region filters, calculated for Samovar, MuTect2, and MosaicHunter variant calls stratified by mosaic allele fraction (MAF) in whole genome sequencing data (WGS). (a) 30X coverage (b) 60X coverage*

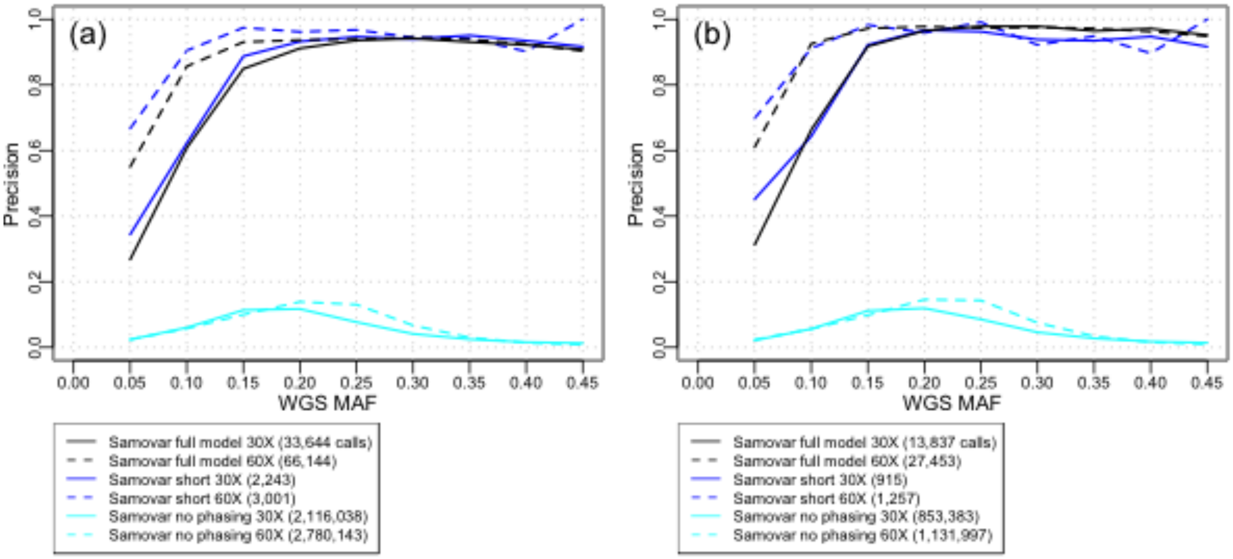

*Figure S4: Precision calculated for variant calls made by Samovar's full model and the "short-only" and "no-phasing" models created for illustration, stratified by mosaic allele fraction (MAF) in whole genome sequencing data (WGS). (a) Autosomes (b) Genomic region not filtered by MosaicHunter or Samovar's region filters*

1. Depth [excluding marked duplicates, QC fail, secondary and supplementary alignments]
2. Fraction of reads phased [HP tag assigned by Long Ranger]
3. Fraction of reads on the more common haplotype [max(number of HP=1 reads, number of HP=2 reads)]
4. MAF
5. MAF of phased reads
6. Number of haplotype-discordant (HD) reads
7. Fraction of phased reads that are HD
8. Fraction of HD reads on the more common haplotype [max(number of HP=1 HD reads, number of HP=2 HD reads)]
9. MAF of HD reads
10. Average base quality of HD reads
11. Average position from the closer end of the alignment on HD reads of the site being classified
12. Average number of soft-clipped bases on HD reads
13. Average number of indels in alignment of HD reads
14. Average value of AS – XS (Lariat alignment scores) of HD reads
15-21. Features 8-14 for the set of phased reads that are not HD
22-26. Features 10-14 for the set of mosaic-allele reads
27-31. Features 10-14 for the set of reference-allele reads
32. "weighted" HD read base quality: sum of HD read base quality / sum of all phased reads base quality
33. "weighted" mosaic-allele read base quality: sum of mosaic-allele read base quality / sum of reference- and mosaic-allele read base quality

***Figure S5: Samovar random forest features***

1. Depth [excluding marked duplicates, QC fail, secondary and supplementary alignments]
2. Fraction of reads phased [computed based on read or its mate overlapping phased variants]
3. Fraction of reads on the more common haplotype [max(number of HP=1 reads, number of HP=2 reads)]
4. MAF
5. MAF of phased reads
6. Number of haplotype-discordant [HD] reads
7. Fraction of phased reads that are HD
8. Fraction of HD reads on the more common haplotype [max(number of HP=1 HD reads, number of HP=2 HD reads)]
9. MAF of HD reads
10. Average base quality of HD reads
11. Average position from the closer end of the alignment on HD reads of the site being classified
12. Average number of soft-clipped bases on HD reads
13. Average number of indels in alignment of HD reads
14-19. Features 8-13 for the set of phased reads that are not HD
20-23. Features 10-13 for the set of mosaic-allele reads
24-27. Features 10-13 for the set of reference-allele reads
28. "weighted" HD read base quality: sum of HD read base quality / sum of all phased reads base quality
29. "weighted" mosaic-allele read base quality: sum of mosaic-allele read base quality / sum of reference- and mosaic-allele read base quality

***Figure S6: Random forest features used in the "short-only" model.***

1. Depth [excluding marked duplicates, QC fail, secondary and supplementary alignments]
2. MAF
3. Average base quality of mosaic-allele reads
4. Average position from the closer end of the alignment on mosaic-allele reads of the site being classified
5. Average number of soft-clipped bases on mosaic-allele reads
6. Average number of indels in alignment of mosaic-allele reads
7-10. Features 3-6 for the set of reference-allele reads
11. "weighted" mosaic-allele read base quality: sum of mosaic-allele read base quality / sum of reference- and mosaic-allele read base quality

***Figure S7: Random forest features used in the "no-phasing" model.***

| **Median depth of mosaic sites** | | | | | | | |
|---|---|---|---|---|---|---|---|
| **13** | # training examples | **1000** | **2000** | **5000** | **10000** | **20000** | |
| (mindepth = 14) | Mosaic >0.5 | 0.9011 | 0.9013 | 0.90304 | 0.90524 | 0.90627 | |
| | Mosaic >0.9 | 0.75662 | 0.7672 | 0.77045 | 0.77511 | 0.77566 | |
| | Het <0.5 | 0.94845 | 0.9491 | 0.94826 | 0.94843 | 0.94747 | |
| | Het <0.9 | 0.98975 | 0.9895 | 0.98991 | 0.99036 | 0.9898 | |
| | Hom <0.5 | 0.95197 | 0.9553 | 0.95669 | 0.95762 | 0.95648 | |
| | Hom <0.9 | 0.99244 | 0.9917 | 0.99265 | 0.99315 | 0.99284 | |
| **25** | # training examples | **1000** | **2000** | **5000** | **10000** | **20000** | **30000** |
| | Mosaic >0.5 | 0.92024 | 0.9185 | 0.92068 | 0.92388 | 0.92355 | 0.92352 |
| | Mosaic >0.9 | 0.80002 | 0.8101 | 0.81813 | 0.81823 | 0.81985 | 0.81635 |
| | Het <0.5 | 0.9598 | 0.9608 | 0.96178 | 0.96032 | 0.95997 | 0.96035 |
| | Het <0.9 | 0.99254 | 0.9912 | 0.99108 | 0.99168 | 0.99144 | 0.99174 |
| | Hom <0.5 | 0.95935 | 0.9636 | 0.96356 | 0.96434 | 0.96431 | 0.96501 |
| | Hom <0.9 | 0.99462 | 0.9938 | 0.99384 | 0.99436 | 0.99528 | 0.99523 |
| **37** | # training examples | **1000** | **2000** | **5000** | **10000** | **20000** | **30000** |
| | Mosaic >0.5 | 0.93072 | 0.933 | 0.93332 | 0.93238 | 0.93283 | 0.93262 |
| | Mosaic >0.9 | 0.82435 | 0.8366 | 0.84427 | 0.84729 | 0.84162 | 0.84557 |
| | Het <0.5 | 0.96298 | 0.9658 | 0.96576 | 0.96782 | 0.96777 | 0.96736 |
| | Het <0.9 | 0.99293 | 0.9924 | 0.99243 | 0.99187 | 0.99252 | 0.9924 |
| | Hom <0.5 | 0.96414 | 0.9685 | 0.96835 | 0.97036 | 0.96997 | 0.97156 |
| | Hom <0.9 | 0.99545 | 0.9951 | 0.99521 | 0.99496 | 0.99551 | 0.99578 |
| **50** | # training examples | **1000** | **2000** | **5000** | **10000** | **20000** | **30000** |
| | Mosaic >0.5 | 0.94165 | 0.9393 | 0.94284 | 0.94151 | 0.94403 | 0.94246 |
| | Mosaic >0.9 | 0.84033 | 0.858 | 0.86627 | 0.86898 | 0.87017 | 0.86613 |
| | Het <0.5 | 0.97008 | 0.9709 | 0.97114 | 0.97159 | 0.97119 | 0.97202 |
| | Het <0.9 | 0.99502 | 0.994 | 0.99365 | 0.99311 | 0.99345 | 0.99341 |
| | Hom <0.5 | 0.96994 | 0.9733 | 0.97322 | 0.97372 | 0.97408 | 0.97418 |
| | Hom <0.9 | 0.99629 | 0.996 | 0.99599 | 0.99566 | 0.99611 | 0.99631 |
| **62** | # training examples | **1000** | **2000** | **5000** | **10000** | **20000** | **30000** |
| | Mosaic >0.5 | 0.95066 | 0.9508 | 0.95103 | 0.94975 | 0.9523 | 0.95019 |
| | Mosaic >0.9 | 0.86295 | 0.8777 | 0.88112 | 0.88244 | 0.88564 | 0.88039 |
| | Het <0.5 | 0.97127 | 0.9725 | 0.97229 | 0.97518 | 0.9736 | 0.97379 |
| | Het <0.9 | 0.99523 | 0.9938 | 0.99411 | 0.99414 | 0.99471 | 0.99444 |
| | Hom <0.5 | 0.96965 | 0.9746 | 0.97586 | 0.9779 | 0.97693 | 0.97807 |
| | Hom <0.9 | 0.99668 | 0.9963 | 0.99674 | 0.99675 | 0.99694 | 0.99707 |

***Table S1: Cross-validation to evaluate the number of training examples and the random forest score threshold.***

1. Minimum depth (excluding marked duplicates, QC fail, secondary and supplementary alignments) [at least 16]
2. Minimum fraction of reads phased [at least 0.5]
3. Minimum fraction of reads on less-prevalent haplotype [at least 0.3]
4. Maximum fraction of reads that have neither reference nor mosaic allele [at most 0.05]
5. Minimum mosaic allele frequency [at least 0.05]
6. Minimum number of haplotype-discordant reads [at least 4]
7. Maximum number of haplotype-discordant reads on the less-prevalent haplotype [at most 0.1]
8. Minimum average position from end of alignment of haplotype-discordant reads [at least 10]

Filters that can be "on" or "off":
1. At least one haplotype-discordant read, one haplotype-concordant read, one reference-allele read and one mosaic-allele read must be aligned in proper pair orientation
2. At least one haplotype-discordant read, one haplotype-concordant read, one reference-allele read and one mosaic-allele read must have an alignment that is not soft-clipped
3. At least one haplotype-discordant read, one haplotype-concordant read, one reference-allele read and one mosaic-allele read must be aligned on the plus and on the minus strand

***Figure S8: preFilter features [default value to pass filter in brackets]***

| | Tumor | | | Normal | | |
|---|---|---|---|---|---|---|
| Case | WES coverage | WGS coverage | CNVNATOR % | WES coverage | WGS coverage | CNVNATOR % |
| 1 | 549 | 45 | 9.3 | 617 | 42 | 8.7 |
| 2 | 504 | 41 | 16.8 | 529 | 41 | 9.3 |
| 3* | 271 | 35 | 23.6 | 255 | 34 | 11.0 |
| 4* | 223 | 34 | 12.4 | 232 | 34 | 11.9 |
| 5* | 207 | 34 | 15.1 | 268 | 35 | 10.9 |
| 6* | 226 | 40 | 11.8 | 223 | 38 | 11.5 |
| 7 | 472 | 35 | 10.3 | 445 | 38 | 8.4 |
| 8* | 330 | 35 | 11.1 | 319 | 34 | 10.9 |
| 9* | 411 | 36 | 16.1 | 346 (Blood)<br>400 (Tissue) | 36 (Blood)<br>36 (Tissue) | 10.8 (Blood)<br>11.0 (Tissue) |
| 10* | 500 | 40 | 11.0 | 392 | 37 | 22.0 |
| 11 | 669 | 37 | 10.5 | 579 | 35 | 10.5 |
| 12* | 618 | 37 | 11.4 | 726 | 37 | 10.9 |
| 13 | 777 | 37 | 20.1 | 681 | 37 | 8.7 |

***Table S2: Cases using reference genome GRCh38 2.1.0 (1, 2, 7, 10, 11) were processed with Long Ranger 2.1.6 and GATK HaplotypeCaller 3.8-0. Samples using reference genome b37 2.1.0 (3, 4, 5, 6, 8, 9, 10, 12) were processed with Long Ranger 2.1.3 and GATK HaplotypeCaller 3.5-0.***

| **Importance** | **Abbreviation** | **Number in Figure S5** |
|---|---|---|
| 0.206699 | weightedMbq | 33 |
| 0.136303 | MAF | 4 |
| 0.115912 | MAF_phased | 5 |
| 0.101952 | weightedCbq | 32 |
| 0.078008 | fracC | 7 |
| 0.075791 | CMAF | 9 |
| 0.065965 | nC | 6 |
| 0.058420 | Mavgbq | 22 |
| 0.050114 | Cavgbq | 10 |
| 0.028026 | NMAF | 16 |
| 0.016379 | Mavgclip | 24 |
| 0.009496 | MavgASXS | 26 |
| 0.008695 | Mavgind | 25 |
| 0.007776 | NavgASXS | 21 |
| 0.006754 | JavgASXS | 31 |
| 0.006130 | CavgASXS | 14 |
| 0.003744 | Cfrach | 8 |
| 0.003665 | Cavgind | 13 |
| 0.003250 | Cavgclip | 12 |
| 0.002759 | Navgbq | 17 |
| 0.002578 | Navgind | 20 |
| 0.002276 | Javgind | 30 |
| 0.001835 | Javgbq | 27 |
| 0.001569 | Javgclip | 29 |
| 0.001236 | fracphased | 2 |
| 0.001149 | depth | 1 |
| 0.000996 | Navgpos | 18 |
| 0.000904 | Mavgpos | 23 |
| 0.000504 | Cavgpos | 11 |
| 0.000476 | Javgpos | 28 |
| 0.000405 | Navgclip | 19 |
| 0.000148 | frach | 3 |
| 0.000086 | Nfrach | 15 |

***Table S3: Samovar model feature importances in simulation experiment***

| | Samovar | | | | | | | | | MuTect2 | | | | | | MosaicHunter | | | | | | | | |
|---|---|---|---|---|---|---|---|---|---|---|---|---|---|---|---|---|---|---|---|---|---|---|---|---|
| | Full Model | | | Short | | | No Phasing | | | Tumor-Only | | | Paired | | | Tumor-Only | | | Paired | | | Trio | | |
| 30X Coverage | Prec | Rec | $F_{0.5}$ | Prec | Rec | $F_{0.5}$ | Prec | Rec | $F_{0.5}$ | Prec | Rec | $F_{0.5}$ | Prec | Rec | $F_{0.5}$ | Prec | Rec | $F_{0.5}$ | Prec | Rec | $F_{0.5}$ | Prec | Rec | $F_{0.5}$ |
| Autosomes | 84.0 | 30.1 | 61.9 | 83.7 | 2.0 | 9.1 | 3.4 | 68.3 | 4.2 | 3.0 | 82.3 | 3.7 | 60.8 | 91.4 | 65.1 | 31.5 | 5.1 | 15.5 | 79.2 | 20.7 | 50.5 | 70.4 | 20.7 | 47.5 |
| Exons | 84.0 | 28.3 | 60.3 | 85.5 | 1.8 | 8.2 | 4.6 | 70.7 | 5.6 | 3.6 | 85.5 | 4.5 | 60.1 | 92.0 | 64.6 | 35.0 | 7.1 | 19.6 | 82.1 | 30.8 | 61.6 | 73.7 | 30.8 | 57.6 |
| Genes | 84.9 | 30.1 | 62.2 | 84.5 | 1.8 | 8.5 | 3.9 | 69.2 | 4.9 | 3.2 | 83.5 | 4.0 | 63.0 | 92.0 | 67.3 | 32.6 | 5.7 | 16.7 | 79.9 | 22.7 | 53.2 | 71.2 | 22.7 | 49.9 |
| Enhancer | 88.5 | 31.0 | 64.6 | 90.9 | 2.2 | 9.9 | 4.4 | 61.8 | 5.4 | 3.9 | 84.6 | 4.9 | 72.9 | 92.3 | 76.1 | 37.8 | 5.9 | 18.1 | 85.5 | 29.5 | 62.0 | 80.2 | 29.5 | 59.7 |
| Promoter | 83.3 | 26.1 | 57.9 | 76.9 | 1.4 | 6.6 | 4.0 | 65.2 | 4.9 | 3.0 | 81.5 | 3.8 | 59.4 | 90.9 | 63.9 | 35.3 | 6.1 | 18.0 | 80.5 | 25.1 | 55.9 | 73.7 | 25.1 | 53.1 |
| Alu | 82.0 | 28.6 | 59.7 | 81.1 | 2.3 | 10.3 | 2.7 | 73.1 | 3.4 | 2.3 | 77.0 | 2.9 | 54.5 | 88.4 | 59.0 | 8.6 | 0.0 | 0.2 | 56.5 | 0.3 | 1.4 | 53.1 | 0.3 | 1.4 |
| RepeatMasker | 84.2 | 29.6 | 61.6 | 82.3 | 2.0 | 9.3 | 2.9 | 67.0 | 3.6 | 2.8 | 80.6 | 3.4 | 58.9 | 90.1 | 63.2 | 20.2 | 0.3 | 1.4 | 72.3 | 1.4 | 6.4 | 61.3 | 1.4 | 6.4 |
| Seg. Dup. | 25.6 | 10.4 | 19.8 | 51.9 | 0.8 | 3.7 | 0.6 | 25.5 | 0.8 | 1.3 | 55.7 | 1.6 | 18.4 | 62.8 | 21.4 | 6.6 | 0.5 | 1.8 | 39.3 | 1.7 | 7.1 | 29.1 | 1.7 | 6.8 |
| 60X Coverage | Prec | Rec | $F_{0.5}$ | Prec | Rec | $F_{0.5}$ | Prec | Rec | $F_{0.5}$ | Prec | Rec | $F_{0.5}$ | | | | Prec | Rec | $F_{0.5}$ | | | | Prec | Rec | $F_{0.5}$ |
| Autosomes | 84.6 | 43.0 | 70.9 | 87.8 | 2.0 | 9.3 | 3.2 | 67.9 | 3.9 | 3.6 | 76.0 | 4.5 | | | | 32.4 | 15.5 | 26.6 | | | | 46.8 | 27.2 | 40.9 |
| Exons | 84.3 | 41.8 | 70.1 | 87.3 | 1.7 | 7.7 | 4.6 | 69.4 | 5.7 | 4.7 | 79.6 | 5.8 | | | | 38.5 | 25.3 | 34.9 | | | | 54.0 | 45.5 | 52.1 |
| Genes | 85.6 | 43.4 | 71.7 | 89.1 | 2.0 | 9.0 | 4.0 | 68.9 | 4.9 | 3.9 | 77.2 | 4.9 | | | | 33.1 | 17.0 | 27.8 | | | | 47.7 | 30.0 | 42.6 |
| Enhancer | 90.8 | 47.8 | 77.0 | 93.3 | 2.2 | 9.9 | 4.4 | 61.1 | 5.3 | 4.8 | 77.9 | 5.9 | | | | 36.9 | 22.7 | 32.8 | | | | 51.6 | 40.0 | 48.8 |
| Promoter | 85.4 | 40.7 | 70.0 | 83.1 | 1.5 | 6.9 | 4.0 | 64.5 | 4.9 | 4.0 | 76.8 | 4.9 | | | | 38.5 | 21.1 | 33.0 | | | | 56.4 | 40.5 | 52.3 |
| Alu | 81.1 | 42.9 | 68.8 | 84.6 | 2.5 | 11.0 | 2.6 | 72.7 | 3.2 | 3.0 | 68.0 | 3.6 | | | | 16.5 | 0.2 | 1.1 | | | | 31.7 | 0.5 | 2.3 |
| RepeatMasker | 84.2 | 42.2 | 70.2 | 87.1 | 2.1 | 9.5 | 2.6 | 66.6 | 3.2 | 3.4 | 74.1 | 4.1 | | | | 24.7 | 1.0 | 4.3 | | | | 38.3 | 1.8 | 7.4 |
| Seg. Dup. | 28.0 | 13.1 | 22.8 | 64.3 | 0.7 | 3.2 | 0.5 | 23.6 | 0.7 | 1.6 | 48.5 | 2.0 | | | | 9.8 | 1.5 | 4.7 | | | | 18.5 | 2.7 | 8.6 |

***Table S4: Precision (Prec), recall (Rec), and $F_{0.5}$ score of each tool for the synthetic mosaic variants inserted by bamsurgeon. This table includes the Samovar "short" and "no-phasing" models, engineered to demonstrate the importance of linked reads for recall and phasing information for precision.***

| | Samovar | | | | | | | | | MuTect2 | | | | | | MosaicHunter | | | | | | | | |
|---|---|---|---|---|---|---|---|---|---|---|---|---|---|---|---|---|---|---|---|---|---|---|---|---|
| | Full Model | | | Short | | | No Phasing | | | Tumor-Only | | | Paired | | | Tumor-Only | | | Paired | | | Trio | | |
| 30X Coverage | Prec | Rec | $F_{0.5}$ | Prec | Rec | $F_{0.5}$ | Prec | Rec | $F_{0.5}$ | Prec | Rec | $F_{0.5}$ | Prec | Rec | $F_{0.5}$ | Prec | Rec | $F_{0.5}$ | Prec | Rec | $F_{0.5}$ | Prec | Rec | $F_{0.5}$ |
| Autosomes | 89.6 | 42.1 | 73.1 | 86.9 | 2.7 | 12.0 | 3.6 | 94.1 | 4.5 | 3.2 | 84.6 | 4.0 | 66.1 | 93.2 | 70.2 | 31.6 | 14.8 | 25.8 | 79.3 | 59.4 | 74.3 | 70.5 | 59.4 | 67.9 |
| Exons | 93.8 | 39.6 | 73.6 | 83.7 | 2.1 | 9.7 | 5.6 | 94.7 | 6.9 | 4.1 | 87.1 | 5.1 | 64.7 | 93.4 | 68.9 | 34.9 | 12.5 | 25.7 | 82.4 | 54.2 | 74.6 | 73.9 | 54.2 | 68.9 |
| Genes | 90.9 | 42.4 | 74.0 | 87.9 | 2.5 | 11.2 | 4.6 | 94.8 | 5.7 | 3.4 | 85.9 | 4.3 | 67.1 | 93.7 | 71.2 | 32.7 | 15.1 | 26.5 | 80.0 | 60.1 | 75.0 | 71.2 | 60.2 | 68.7 |
| Enhancer | 94.2 | 42.0 | 75.4 | 91.7 | 2.6 | 11.7 | 5.5 | 96.1 | 6.8 | 4.0 | 85.3 | 5.0 | 70.1 | 93.6 | 73.8 | 36.4 | 11.3 | 25.2 | 85.9 | 58.7 | 78.6 | 80.1 | 58.7 | 74.6 |
| Promoter | 91.4 | 36.3 | 70.1 | 76.7 | 1.8 | 8.0 | 4.6 | 93.9 | 5.7 | 3.2 | 84.6 | 4.0 | 60.5 | 92.4 | 65.0 | 35.4 | 11.6 | 25.1 | 80.7 | 48.7 | 71.3 | 74.2 | 48.7 | 67.1 |
| Alu | 30.2 | 18.6 | 26.9 | 25.0 | 1.4 | 5.8 | 0.9 | 60.5 | 1.1 | 1.4 | 53.1 | 1.8 | 27.1 | 61.4 | 30.5 | 9.7 | 4.3 | 7.7 | 52.6 | 28.6 | 45.0 | 50.0 | 28.6 | 43.5 |
| RepeatMasker | 68.5 | 33.1 | 56.4 | 73.7 | 2.3 | 10.3 | 0.7 | 73.9 | 0.8 | 2.6 | 71.5 | 3.2 | 45.6 | 75.8 | 49.6 | 24.4 | 10.1 | 19.0 | 75.5 | 45.0 | 66.5 | 65.3 | 45.0 | 59.9 |
| Seg. Dup. | 6.8 | 4.4 | 6.1 | 28.6 | 0.6 | 2.9 | 0.1 | 17.0 | 0.2 | 0.8 | 42.6 | 1.0 | 11.1 | 39.6 | 13.0 | 7.8 | 4.1 | 6.6 | 37.6 | 11.9 | 26.3 | 27.7 | 12.3 | 22.1 |
| 60X Coverage | Prec | Rec | $F_{0.5}$ | Prec | Rec | $F_{0.5}$ | Prec | Rec | $F_{0.5}$ | Prec | Rec | $F_{0.5}$ | | | | Prec | Rec | $F_{0.5}$ | | | | Prec | Rec | $F_{0.5}$ |
| Autosomes | 89.7 | 60.3 | 81.7 | 89.3 | 2.7 | 12.2 | 3.4 | 94.0 | 4.2 | 4.0 | 78.7 | 4.9 | | | | 32.4 | 44.5 | 34.2 | | | | 46.8 | 78.3 | 50.9 |
| Exons | 91.7 | 58.5 | 82.4 | 89.2 | 2.1 | 9.5 | 5.9 | 95.0 | 7.3 | 5.4 | 81.8 | 6.6 | | | | 38.6 | 44.7 | 39.7 | | | | 54.0 | 80.6 | 57.9 |
| Genes | 90.8 | 60.8 | 82.6 | 91.3 | 2.7 | 12.0 | 5.0 | 95.1 | 6.1 | 4.3 | 79.8 | 5.2 | | | | 33.1 | 45.0 | 34.9 | | | | 47.6 | 79.4 | 51.8 |
| Enhancer | 94.5 | 63.3 | 86.0 | 94.4 | 2.8 | 12.7 | 5.8 | 94.8 | 7.1 | 4.9 | 79.6 | 6.1 | | | | 36.5 | 45.1 | 38.0 | | | | 51.1 | 79.6 | 55.1 |
| Promoter | 91.2 | 57.3 | 81.5 | 92.2 | 2.3 | 10.5 | 4.9 | 94.7 | 6.1 | 4.2 | 78.2 | 5.2 | | | | 38.6 | 40.5 | 39.0 | | | | 56.5 | 78.1 | 59.9 |
| Alu | 45.9 | 30.0 | 41.5 | 57.1 | 3.1 | 12.7 | 0.7 | 53.1 | 0.9 | 2.1 | 56.2 | 2.6 | | | | 17.0 | 20.8 | 17.6 | | | | 32.4 | 43.8 | 34.2 |
| RepeatMasker | 73.6 | 43.9 | 64.8 | 82.7 | 2.5 | 11.0 | 0.4 | 69.4 | 0.5 | 3.2 | 62.8 | 3.9 | | | | 27.5 | 31.2 | 28.1 | | | | 41.5 | 54.9 | 43.6 |
| Seg. Dup. | 9.2 | 6.0 | 8.3 | 15.4 | 0.4 | 1.6 | 0.1 | 13.3 | 0.1 | 1.0 | 32.9 | 1.3 | | | | 10.1 | 10.3 | 10.2 | | | | 18.8 | 18.5 | 18.8 |

***Table S5: Precision (Prec), recall (Rec), and $F_{0.5}$ score of each tool for the synthetic mosaic variants inserted by bamsurgeon in the region of the genome not filtered out by MosaicHunter or Samovar. This table includes the Samovar "short" and "no-phasing" models, engineered to demonstrate the importance of linked reads for recall and phasing information for precision.***

| Importance | Abbreviation | Number in Figure S6 |
|---|---|---|
| 0.20520657 | weightedMbq | 33 |
| 0.16749462 | MAF | 4 |
| 0.13410861 | weightedCbq | 32 |
| 0.08401046 | fracC | 7 |
| 0.07369292 | MAF_phased | 5 |
| 0.06894606 | Mavgbq | 22 |
| 0.06203258 | nC | 6 |
| 0.02828793 | Cfrach | 8 |
| 0.02817725 | Mavgclip | 24 |
| 0.02439597 | Cavgbq | 10 |
| 0.01783998 | MavgASXS | 26 |
| 0.01466872 | JavgASXS | 31 |
| 0.01386411 | CMAF | 9 |
| 0.01311324 | CavgASXS | 14 |
| 0.01274496 | NMAF | 16 |
| 0.00876721 | NavgASXS | 21 |
| 0.008098 | Mavgind | 25 |
| 0.00757443 | fracphased | 2 |
| 0.00331775 | Navgind | 20 |
| 0.00318227 | Javgind | 30 |
| 0.00307518 | Cavgind | 13 |
| 0.00297741 | Mavgpos | 23 |
| 0.00291929 | Cavgpos | 11 |
| 0.00285171 | Javgbq | 27 |
| 0.00209531 | Javgclip | 29 |
| 0.00169362 | Navgbq | 17 |
| 0.00123443 | Javgpos | 28 |
| 0.00070641 | Navgclip | 19 |
| 0.00070569 | Nfrach | 15 |
| 0.00069773 | Navgpos | 18 |
| 0.00069729 | depth | 1 |
| 0.00052806 | frach | 3 |
| 0.00029423 | Cavgclip | 12 |

***Table S6: Short-read phasing model feature importances in simulation experiment***

| Importance | Abbreviation | Number in Figure |
|---:|---:|---:|
| 0.42298002 | weightedMbq | 11 |
| 0.29348068 | MAF | 2 |
| 0.14507064 | Mavgbq | 3 |
| 0.07905771 | Mavgclip | 5 |
| 0.03207496 | Mavgind | 6 |
| 0.00746506 | Javgind | 10 |
| 0.00448625 | Javgpos | 8 |
| 0.00438944 | Javgbq | 7 |
| 0.00403807 | depth | 1 |
| 0.0038253 | Mavgpos | 4 |
| 0.00313186 | Javgclip | 9 |

***Table S7: No-phasing model feature importances in simulation experiment***

| Case | Diagnosis | Sex* |
|---|---|---|
| 1 | Indeterminate, most consistent with oligodendroglioma | M |
| 2 | Pilocytic astrocytoma | M |
| 3 | Medulloblastoma, WHO grade IV,<br>most consistent with non-WNT, non-SHH subgroup | M |
| 4 | Pilocytic astrocytoma | F |
| 5 | Glioblastoma (recurrence) | F |
| 6 | Pilocytic astrocytoma | F |
| 7 | Ewing-like sarcoma | M |
| 8 | Ganglioglioma, WHO grade 1 | M |
| 9 | Diffuse Midline Glioma, H3 K27M-mutant, WHO grade IV | M |
| 10 | Indeterminate, high grade glioma/astrocytoma | M |
| 11 | Ganglioglioma, WHO grade 1 | F |
| 12 | Glioma (low grade) | M |
| 13 | Clival chordoma | F |

***Table S8: Metadata for each case. * Sex determined from alignments to Y-chromosome***

| Case | Calls | Sensitivity |
| --- | --- | --- |
| 1 | 58 | 0.70 |
| 2 | 85 | 0.61 |
| 3 | 70 | 0.57 |
| 4 | 73 | 0.68 |
| 5 | 51 | 0.58 |
| 6 | 73 | 0.58 |
| 7 | 61 | 0.63 |
| 8 | 43 | 0.62 |
| 9 | 39 | 0.48 |
| 10 | 50 | 0.45 |
| 11 | 30 | 0.73 |
| 12 | 59 | 0.78 |
| 13 | 70 | 0.88 |
| Total | 762 | |

***Table S9: Samovar analysis of normal WGS dataset for pediatric cancer cases. Number of calls shown is for the WES capture region, and validation performed as described in main text.***

**Note S1. Samovar requirements .**
Samovar is implemented in Python 3 (also compatible with Python 2). It uses several libraries, including pyfaidx, scikit-learn, simplesam, and fisher. As input, Samovar requires the alignment (BAM) and variant (VCF) files produced by 10x Genomics' Long Ranger pipeline. Long Ranger processes the raw Illumina reads and performs linked read-aware alignment with Lariat [20], small variant calling with Freebayes [47] or GATK [48], structural variant calling, and haplotype assembly. Specifically, Samovar requires that the BAM have the HP (molecule haplotype), AS (Lariat best alignment score), and XS (Lariat second-best alignment score) extra fields, and requires that the VCF have the FILTER column and GT field. Information on the BAM file tags and phased VCF file are available at 10x Genomics.

**Note S2. Computational performance .**
Running time for each tool was listed for the 30X simulation experiment described in the text. Samovar was run on a single machine with 48 cores for the "filter" step and 4 cores for other parallelizable steps, with pypy when possible. Maximum memory usage was 19.2 GB, and the filter step reported 4200% CPU usage when allocated 48 cores. MosaicHunter and MuTect2 were run on a cluster in a scatter-gather format where each chromosome was computed independently and the results were merged. MosaicHunter does not offer parallelism options, although slightly greater than 100% average CPU usage was seen. On chromosome 1, paired mode used maximum 25.6 GB memory; tumor-only mode used 25.3 GB; trio mode used 25.0 GB. MuTect2 was run with 48 cores for the "native pair HMM," although only 600% CPU usage was seen on average. On chromosome 1, paired mode used maximum 5.7 GB memory; tumor-only mode used 5.5 GB.

**Note S3. Command line arguments .**

**MosaicHunter** Version 1.1
The tumor-only, paired, and trio configuration file templates provided with the software distribution

were used, containing default parameters.

```
java -jar mosaichunter.jar -C [configuration file] -P output_dir=[output directory]
```

**MuTect2 - Paired Mode** Version 4.0.12.0

```
gatk Mutect2 -R [reference genome] -I [tumor BAM file] -tumor [tumor sample name] -I
[normal BAM file] -normal [normal sample name] -O [MuTect2 VCF file]
--native-pair-hmm-threads 48
gatk FilterMutectCalls -V [MuTect2 VCF file] -O [Filtered VCF file]
grep -v "multiallelic" [Filtered VCF file] | grep -v "0/1/2" | vcftools --vcf -
--remove-indels --remove-filtered-all --recode --recode-INFO-all --out [Final MuTect2
VCF file]
```

**MuTect2 - Tumor-only mode** Version 4.0.12.0

```
gatk Mutect2 -R [reference genome] -I [tumor BAM file] -tumor [tumor sample name] -O
[MuTect2 VCF file] --native-pair-hmm-threads 48
gatk FilterMutectCalls -V [MuTect2 VCF file] -O [Filtered VCF file]
grep -v "multiallelic" [Filtered VCF file] | grep -v "0/1/2" | vcftools --vcf
- --remove-indels --remove-filtered-all --recode --recode-INFO-all --out [Final MuTect2
VCF file]
```

**Samovar**

```
samovar generateVarfile --out out.varfile --vcf [Sample VCF] --fai [Reference genome
FAI]
samovar simulate --bam [Sample BAM] --varfile [Sample VCF] --het --max 15000 --nproc
4 > het.features.tsv
samovar simulate --bam [Sample BAM] --varfile [Sample VCF] --hom --max 15000 --nproc
4 > hom.features.tsv
samovar simulate --bam [Sample BAM] --varfile out.varfile --simulate --nproc 4
> mosaic.features.tsv
samovar train --mosaic mosaic.features.tsv --het het.features.tsv --hom hom.features.tsv
samovar preFilter --bam [Sample BAM --nproc 48 > vectors.txt 2> intervalsComplete.txt
samovar classify --clf clf.pkl --vectors vectors.txt > predictions.tsv
bedtools intersect -v -a predictions.tsv -b [Samovar repeat BED file] | bedtools
intersect -v -a stdin -b [CNVNATOR BED file] > regionfiltered.tsv
samovar postFilter --bam [Sample BAM] --bed regionfiltered.tsv --ref [Reference genome]
--vcfavoid [Sample VCF] --nproc 4 --p 0.005 > samovar.vcf
```

**Note S4. Genomic regions and filters** .

In Table 1 and Figure 3 Samovar and MosaicHunter use their respective default filters but we have treated the tools as though they are interrogating roughly the same portion of the genome. Table S5 and Figure S3 attempt to normalize the differences by reporting just those sites that pass both tools' filters. In GRCh38, this is 32.8% of the autosomal sequence, containing MosaicHunter's simple sequence repeat filter and repetitive region bed files, and Samovar's simple sequence repeat filter, as well as any CNV regions identified by CNVNATOR.

### Supplementary File 1

| Case in Table 2, S2 | NCH Case ID | Library Type(s) | 10X Prep Identifier | Processing Number |
|---|---|---|---|---|
| 11 | JL_17014_M17-2600 | 10XG WGS and WES | 10XTEST12 | 1 |
| 11 | JL_17014_M17-2600 | 10XG WGS and WES | 10XTEST12 | 2 |
| 12 | JL_17015_M17-2670 | 10XG WGS and WES | 10XTEST12 | 3 |
| 12 | JL_17015_M17-2670 | 10XG WGS and WES | 10XTEST12 | 4 |
| 13 | JL_17016_M17-2796 | 10XG WGS and WES | 10XTEST12 | 5 |
| 13 | JL_17016_M17-2796 | 10XG WGS and WES | 10XTEST12 | 6 |
| 10 | JL_17013_M17-2121 | 10XG WGS and WES | 10XTEST10 | 1 |
| 10 | JL_17013_M17-2121 | 10XG WGS and WES | 10XTEST10 | 2 |
| 7 | JL_17009_M17-2428 | 10XG WGS and WES | 10XTEST10 | 3 |
| 7 | JL_17009_M17-2428 | 10XG WGS and WES | 10XTEST10 | 4 |
| 8 | JL_17011_M17-2425 | 10XG WGS and WES | 10XTEST10 | 5 |
| 8 | JL_17011_M17-2425 | 10XG WGS and WES | 10XTEST10 | 6 |
| 9 | JL_17012_M17-2423 | 10XG WGS and WES | 10XTEST10 | 7 |
| 9 | JL_17012_M17-2423 | 10XG WGS and WES | 10XTEST10 | 8 |
| 9 | JL_17012_M17-2423 | 10XG WGS and WES | 10XTEST11 | 1 |
| 1 | JL_17001_M17-1537 | 10XG WGS and WES | 10XTEST4 | 1 |
| 1 | JL_17001_M17-1537 | 10XG WGS and WES | 10XTEST4 | 2 |
| 2 | JL_17002_M17-1836 | 10XG WGS and WES | 10XTEST7 | 3 |
| 2 | JL_17002_M17-1836 | 10XG WGS and WES | 10XTEST7 | 4 |
| 3 | JL_17003_M17-1870 | 10XG WGS and WES | 10XTEST8 | 1 |
| 3 | JL_17003_M17-1870 | 10XG WGS and WES | 10XTEST8 | 2 |
| 4 | JL_17004_M17-1862 | 10XG WGS and WES | 10XTEST8 | 3 |
| 4 | JL_17004_M17-1862 | 10XG WGS and WES | 10XTEST8 | 4 |
| 5 | JL_17005_M17-1890 | 10XG WGS and WES | 10XTEST8 | 5 |
| 5 | JL_17005_M17-1890 | 10XG WGS and WES | 10XTEST8 | 6 |
| 6 | JL_17006_M17-1974 | 10XG WGS and WES | 10XTEST8 | 7 |
| 6 | JL_17006_M17-1974 | 10XG WGS and WES | 10XTEST8 | 8 |

| 10XG Sequencing Run Identifier | 10XG Sequencing Run Concentration | No. of Lanes |
|---|---|---|
| IGM-HiSeq4000-117-0194B; 9-29-17; 151,8,151 | 3nM | 1.33 |
| IGM-HiSeq4000-117-0194B; 9-29-17; 151,8,151 | 3nM | 1.33 |
| IGM-HiSeq4000-117-0194B; 9-29-17; 151,8,151 | 3nM | 1.33 |
| IGM-HiSeq4000-117-0194B; 9-29-17; 151,8,151 | 3nM | 1.33 |
| IGM-HiSeq4000-117-0194B; 9-29-17; 151,8,151 | 3nM | 1.33 |
| IGM-HiSeq4000-117-0194B; 9-29-17; 151,8,151 | 3nM | 1.33 |
| BGC-HiSeq4000-102-B; 8-11-17; 151,8,151 | 3nM | 1.33 |
| BGC-HiSeq4000-102-B; 8-11-17; 151,8,151 | 3nM | 1.33 |
| BGC-HiSeq4000-102-B; 8-11-17; 151,8,151 | 3nM | 1.33 |
| BGC-HiSeq4000-102-B; 8-11-17; 151,8,151 | 3nM | 1.33 |
| BGC-HiSeq4000-102-B; 8-11-17; 151,8,151 | 3nM | 1.33 |
| BGC-HiSeq4000-102-B; 8-11-17; 151,8,151 | 3nM | 1.33 |
| IGM-HiSeq4000-106-A; 8-21-17; 151,8,151 | 3nM | 1.25 |
| IGM-HiSeq4000-106-A; 8-21-17; 151,8,151 | 3nM | 1.25 |
| IGM-HiSeq4000-106-A; 8-21-17; 151,8,151 | 3nM | 1.25 |
| BGC-HiSeq4000-075; 5-5-17; 151,8,8,151 | 3nM | 1.5 |
| BGC-HiSeq4000-075; 5-5-17; 151,8,8,151 | 3nM | 1.5 |
| BGC-HiSeq4000-084; 6-2-17; 151,8,151 | 3nM | 1.5 |
| BGC-HiSeq4000-084; 6-2-17; 151,8,151 | 3nM | 1.5 |
| BGC-HiSeq4000-088; 6-19-17; 151,8,151 | 3nM | 1.33 |
| BGC-HiSeq4000-088; 6-19-17; 151,8,151 | 3nM | 1.33 |
| BGC-HiSeq4000-088; 6-19-17; 151,8,151 | 3nM | 1.33 |
| BGC-HiSeq4000-088; 6-19-17; 151,8,151 | 3nM | 1.33 |
| BGC-HiSeq4000-088; 6-19-17; 151,8,151 | 3nM | 1.33 |
| BGC-HiSeq4000-088; 6-19-17; 151,8,151 | 3nM | 1.33 |
| BGC-HiSeq4000-089; 6-21-17; 151,8,8,151 | 3nM | 1.5 |
| BGC-HiSeq4000-089; 6-21-17; 151,8,8,151 | 3nM | 1.5 |

| Sample Name | Extraction Method |
|---|---|
| 17-0615-03_H3950_Tumor | AllPrep DNA |
| 17-0617-03_H3390_BloodNormal | Puregene |
| 17-0618-03_H3949_Tumor | AllPrep DNA |
| 17-0620-03_H3549_BloodNormal | Puregene |
| 17-0621-03_H3948_Tumor | AllPrep DNA |
| 17-0623-03_H3633_BloodNormal | Puregene |
| 17-0469-03_H3182_Tumor | AllPrep DNA |
| 17-0471-03_H2397_BloodNormal | Puregene |
| 17-0472-03_H3184_Tumor | AllPrep DNA |
| 17-0474-03_H3086_BloodNormal | Puregene |
| 17-0475-03_H3186_Tumor | AllPrep DNA |
| 17-0477-03_H3108_BloodNormal | Puregene |
| 17-0478-03_H3187_Tumor | AllPrep DNA |
| 17-0480-03_H3188_TissueNormal | AllPrep DNA |
| 17-0482-03_H3087_BloodNormal | Puregene |
| 17-0019-03_H1575_Normal | AllPrep DNA |
| 17-0020-03_H1576_Tumor | QIAamp DNA |
| 17-0094-03_H2090_Tumor | AllPrep DNA |
| 17-0095-03_H2091_Normal | QIAamp DNA |
| 17-0216-03_H2089_Tumor | AllPrep DNA |
| 17-0217-03_H2217_Normal | QIAamp DNA |
| 17-0219-03_H2218_Tumor | AllPrep DNA |
| 17-0220-03_H2219_Normal | QIAamp DNA |
| 17-0222-03_H2220_Tumor | AllPrep DNA |
| 17-0223-03_H2221_Normal | QIAamp DNA |
| 17-0303-03_H2315_Tumor | AllPrep DNA |
| 17-0304-03_H2316_Normal | QIAamp DNA |

| Average concentration after dilutions (ng/uL) | Input for GEM Generation (ng) | Date of Post GEM Incubation | Peak after GEM Generation and Barcoding (bp) | Average Size after GEM Generation and Barcoding (bp) | Date of Library Construction |
|---|---|---|---|---|---|
| 1.004 | 1.254 | 9/21/17 | 520 | 828 | 9/22/17 |
| 0.914 | 1.142 | 9/21/17 | 591 | 859 | 9/22/17 |
| 1.000 | 1.250 | 9/21/17 | 538 | 851 | 9/22/17 |
| 0.830 | 1.038 | 9/21/17 | 550 | 836 | 9/22/17 |
| 1.050 | 1.313 | 9/21/17 | 492 | 821 | 9/22/17 |
| 0.972 | 1.214 | 9/21/17 | 505 | 809 | 9/22/17 |
| 0.993 | 2.481 | 8/8/17 | 380 | 778 | 8/9/17 |
| 1.020 | 2.550 | 8/8/17 | 557 | 859 | 8/9/17 |
| 1.055 | 2.638 | 8/8/17 | 536 | 844 | 8/9/17 |
| 1.075 | 2.688 | 8/8/17 | 510 | 836 | 8/9/17 |
| 1.080 | 2.700 | 8/8/17 | 533 | 842 | 8/9/17 |
| 0.948 | 2.370 | 8/8/17 | 505 | 835 | 8/9/17 |
| 1.085 | 2.713 | 8/8/17 | 450 | 789 | 8/9/17 |
| 1.050 | 2.625 | 8/8/17 | 479 | 813 | 8/9/17 |
| 0.999 | 1.248 | 8/10/17 | 530 | 841 | 8/14/17 |
| 1.095 | 1.369 | 5/2/17 | 511 | 850 | 5/3/17 |
| 1.170 | 1.463 | 5/2/17 | 578 | 867 | 5/3/17 |
| 1.195 | 1.494 | 5/24/17 | 514 | 875 | 5/25/17 |
| 0.970 | 1.213 | 5/24/17 | 559 | 898 | 5/25/17 |
| 1.064 | 1.329 | 6/12/17 | 605 | 839 | 6/13/17 |
| 1.070 | 1.338 | 6/12/17 | 598 | 814 | 6/13/17 |
| 1.090 | 1.363 | 6/12/17 | 567 | 857 | 6/13/17 |
| 1.070 | 1.338 | 6/12/17 | 579 | 847 | 6/13/17 |
| 1.040 | 1.300 | 6/12/17 | 568 | 892 | 6/13/17 |
| 0.995 | 1.244 | 6/12/17 | 434 | 830 | 6/13/17 |
| 1.060 | 1.325 | 6/12/17 | 474 | 825 | 6/13/17 |
| 1.055 | 1.319 | 6/12/17 | 465 | 841 | 6/13/17 |

| Index Number | Index Sequence1 | Index Sequence2 | Index Sequence3 | Index Sequence4 |
|---|---|---|---|---|
| SI-GA-G5 | GAGCAAGA | TCTGTGAT | CGCAGTTC | ATATCCCG |
| SI-GA-G6 | CTGACGCG | GGTCGTAC | TCCTTCTT | AAAGAAGA |
| SI-GA-G7 | GGTATGCA | CTCGAAAT | ACACCTTC | TAGTGCGG |
| SI-GA-G8 | TATGAGCT | CCGATAGC | ATACCCAA | GGCTGTTG |
| SI-GA-G9 | TAGGACGT | ATCCCACA | GGAATGTC | CCTTGTAG |
| SI-GA-G10 | TCGCCAGC | AATGTTAG | CGATAGCT | GTCAGCTA |
| SI-GA-F5 | GACTACGT | CTAGCGAG | TCTATATC | AGGCGTCA |
| SI-GA-F6 | CGGAGCAC | GACCTATT | ACTTAGGA | TTAGCTCG |
| SI-GA-F7 | CGTGCAGA | AACAAGAT | TCGCTTCG | GTATGCTC |
| SI-GA-F8 | CATGAACA | TCACTCGC | AGCTGGAT | GTGACTTG |
| SI-GA-F9 | CAAGCTCC | GTTCACTG | TCGTGAAA | AGCATGGT |
| SI-GA-F10 | GCTTGGCT | AAACAAAC | CGGGCTTA | TTCATCGG |
| SI-GA-F11 | GCGAGAGT | TACGTTCA | AGTCCCAC | CTATAGTG |
| SI-GA-F12 | TGATGCAT | GCTACTGA | CACCTGCC | ATGGAATG |
| SI-GA-G1 | ATGAATCT | GATCTCAG | CCAGGAGC | TGCTCGTA |
| SI-GA-B6 | CGTTAATC | GCCACGCT | TTACTCAG | AAGGGTGA |
| SI-GA-B7 | AAACCTCA | GCCTTGGT | CTGGACTC | TGTAGAAG |
| SI-GA-D9 | AGGAGATG | GATGTGGT | CTACATCC | TCCTCCAA |
| SI-GA-D10 | CAATACCC | TGTCTATG | ACCACGAA | GTGGGTGT |
| SI-GA-E3 | AGGTATTG | CTCCTAGT | TCAAGGCC | GATGCCAA |
| SI-GA-E4 | TTCGCCCT | GGATGGGC | AATCAATG | CCGATTAA |
| SI-GA-E5 | CATTAGCG | TTCGCTGA | ACAAGAAT | GGGCTCTC |
| SI-GA-E6 | CTGCGGCT | GACTCAAA | AGAAACTC | TCTGTTGG |
| SI-GA-E7 | CACGCCTT | GTATATAG | TCTCGGGC | AGGATACA |
| SI-GA-E8 | ATAGTTAC | TGCTGAGT | CCTACGTA | GAGCACCG |
| SI-GA-E9 | TTGTTTCC | GGAGGAGG | CCTAACAA | AACCCGTT |
| SI-GA-E10 | AAATGTGC | GGGCAAAT | TCTATCCG | CTCGCGTA |

| Peak after Library Construction | Average Size of Insert from Tape Station (bp) | Qubit ng/uL | Total Yield after QC | qPCR of 10XG based on 550bp size |
|---|---|---|---|---|
| 535 | 743 | 22.8 | 433.2 | 33.4 |
| 591 | 803 | 24.4 | 463.6 | 33.1 |
| 532 | 757 | 29.1 | 552.9 | 50.2 |
| 550 | 771 | 26.3 | 499.7 | 48.3 |
| 525 | 733 | 26.2 | 497.8 | 54.5 |
| 524 | 730 | 32.0 | 608.0 | 60.2 |
| 507 | 743 | 43.6 | 827.5 | 100.1 |
| 553 | 715 | 24.4 | 462.7 | 49.7 |
| 574 | 778 | 32.6 | 619.4 | 86.0 |
| 583 | 781 | 39.0 | 741.0 | 85.4 |
| 556 | 780 | 43.4 | 823.7 | 84.8 |
| 582 | 778 | 45.2 | 857.9 | 87.7 |
| 502 | 702 | 41.7 | 791.4 | 94.9 |
| 551 | 742 | 44.1 | 837.9 | 88.9 |
| 601 | 737 | 29.5 | 559.6 | 32.0 |
| 590 | 741 | 28.7 | 545.3 | 38.5 |
| 611 | 760 | 15.8 | 300.2 | 7.5 |
| 543 | 698 | 25.4 | 456.3 | 31.8 |
| 530 | 691 | 33.3 | 599.4 | 48.6 |
| 540 | 683 | 28.3 | 537.7 | 49.7 |
| 570 | 726 | 34.7 | 658.4 | 48.4 |
| 598 | 729 | 36.5 | 692.6 | 43.1 |
| 551 | 708 | 38.2 | 725.8 | 44.0 |
| 592 | 745 | 32.1 | 609.9 | 28.8 |
| 543 | 701 | 37.3 | 707.8 | 52.8 |
| 562 | 712 | 38.8 | 737.2 | 47.7 |
| 550 | 699 | 40.5 | 768.6 | 55.0 |

| 10XG Exome Sequencing Run Identifier | 10XG Exome Sequencing Run Concentration | No. of Lanes |
|---|---|---|
| IGM-HiSeq4000-118-0194A; 10-5-17; 151,8,8,151 | 3nM | 0.66 |
| IGM-HiSeq4000-118-0194A; 10-5-17; 151,8,8,151 | 3nM | 0.66 |
| IGM-HiSeq4000-118-0194A; 10-5-17; 151,8,8,151 | 3nM | 0.66 |
| IGM-HiSeq4000-118-0194A; 10-5-17; 151,8,8,151 | 3nM | 0.66 |
| IGM-HiSeq4000-118-0194A; 10-5-17; 151,8,8,151 | 3nM | 0.66 |
| IGM-HiSeq4000-118-0194A; 10-5-17; 151,8,8,151 | 3nM | 0.66 |
| IGM-HiSeq4000-107-1B; 8-24-17; 151,8,8,151 | 3nM | 0.33 |
| IGM-HiSeq4000-107-1B; 8-24-17; 151,8,8,151 | 3nM | 0.33 |
| IGM-HiSeq4000-107-1B; 8-24-17; 151,8,8,151 | 3nM | 0.33 |
| IGM-HiSeq4000-107-1B; 8-24-17; 151,8,8,151 | 3nM | 0.33 |
| IGM-HiSeq4000-107-1B; 8-24-17; 151,8,8,151 | 3nM | 0.33 |
| IGM-HiSeq4000-107-1B; 8-24-17; 151,8,8,151 | 3nM | 0.33 |
| IGM-HiSeq4000-107-1B; 8-24-17; 151,8,8,151 | 3nM | 0.33 |
| IGM-HiSeq4000-107-1B; 8-24-17; 151,8,8,151 | 3nM | 0.33 |
| IGM-HiSeq4000-107-1B; 8-24-17; 151,8,8,151 | 3nM | 0.33 |
| BGC-HiSeq4000-075; 5-5-17; 151,8,8,151 | 3nM | 0.5 |
| BGC-HiSeq4000-075; 5-5-17; 151,8,8,151 | 3nM | 0.5 |
| BGC-HiSeq4000-084; 6-2-17; 151,8,151 | 3nM | 0.5 |
| BGC-HiSeq4000-084; 6-2-17; 151,8,151 | 3nM | 0.5 |
| BGC-HiSeq4000-089; 6-21-17; 151,8,8,151 | 3nM | 0.25 |
| BGC-HiSeq4000-089; 6-21-17; 151,8,8,151 | 3nM | 0.25 |
| BGC-HiSeq4000-089; 6-21-17; 151,8,8,151 | 3nM | 0.25 |
| BGC-HiSeq4000-089; 6-21-17; 151,8,8,151 | 3nM | 0.25 |
| BGC-HiSeq4000-089; 6-21-17; 151,8,8,151 | 3nM | 0.25 |
| BGC-HiSeq4000-089; 6-21-17; 151,8,8,151 | 3nM | 0.25 |
| BGC-HiSeq4000-089; 6-21-17; 151,8,8,151 | 3nM | 0.25 |
| BGC-HiSeq4000-089; 6-21-17; 151,8,8,151 | 3nM | 0.25 |

| Sample Identifer for 10XG Exome Capture | Volume (uL) Remove 250ng for Exome IDT Capture | DATE of Exome Capture | Number of Cycles | No. of rxn for Amp PCR |
|---|---|---|---|---|
| 17-0615-04_H3950_Tumor | 11.0 | 9/26/17 | 9 | 2 |
| 17-0617-04_H3390_BloodNormal | 10.2 | | | 6 Sample |
| 17-0618-04_H3949_Tumor | 8.6 | | | |
| 17-0620-04_H3549_BloodNormal | 9.5 | | | |
| 17-0621-04_H3948_Tumor | 9.5 | | | |
| 17-0623-04_H3633_BloodNormal | 7.8 | | | |
| 17-0469-04_H3182_Tumor | 5.7 | 8/22/17 | | |
| 17-0471-04_H2397_BloodNormal | 10.3 | | | 9 Sample |
| 17-0472-04_H3184_Tumor | 7.7 | | | |
| 17-0474-04_H3086_BloodNormal | 6.4 | | | |
| 17-0475-04_H3186_Tumor | 5.8 | | | |
| 17-0477-04_H3108_BloodNormal | 5.5 | | | |
| 17-0478-04_H3187_Tumor | 6.0 | | | |
| 17-0480-04_H3188_TissueNormal | 5.7 | | | |
| 17-0482-04_H3087_BloodNormal | 8.5 | | | |
| 17-0019-04_H1575_Normal | 8.7 | 5/4/17 | 12.0 | 2.0 |
| 17-0020-04_H1576_Tumor | 15.8 | | | 2 Sample |
| 17-0094-04_H2090_Tumor | 9.9 | 5/26/17 | 12 | 2 |
| 17-0095-04_H2091_Normal | 7.5 | | | 2 Sample |
| 17-0216-04_H2089_Tumor | 8.8 | 6/14/17 | 9 | 2 |
| 17-0217-04_H2217_Normal | 7.2 | | | 8 Sample |
| 17-0219-04_H2218_Tumor | 6.9 | | | |
| 17-0220-04_H2219_Normal | 6.5 | | | |
| 17-0222-04_H2220_Tumor | 7.8 | | | |
| 17-0223-04_H2221_Normal | 6.7 | | | |
| 17-0303-04_H2315_Tumor | 6.4 | | | |
| 17-0304-04_H2316_Normal | 6.2 | | | |

| Volume used for each rxn Amp PCR | Peak after 10XG-IDTEXome | Average size after 10XG-IDTEXome | qPCR of 10XG Exome based on 550bp size |
|---|---|---|---|
| 4 | 600 | 739 | 31.6 |

s pooled together

| | | | |
|---|---|---|---|
| | | | 20.6 |

s Pooled together

| | | | |
|---|---|---|---|
| 4.0 | 509.0 | 683.0 | 27.5 |

s pooled together

| | | | |
|---|---|---|---|
| 4 | 560 | 688 | 22.4 |

s pooled together

| | | | |
|---|---|---|---|
| 4 | 509 | 638 | 29.6 |

s pooled together
